# Multiparametric microenvironment sensing via distinct molecular equilibria in a single cyanine dye

**DOI:** 10.64898/2026.08.29.747692

**Authors:** Sujata Bais, Soren Westrey, Cecilia Samaniego López, M. Verónica Rivas, Carla C. Spagnuolo, Saumya Saurabh

## Abstract

Reading both physical and chemical properties of a microenvironment from a single fluorophore remains a challenge. Here we demonstrate that two coexisting molecular equilibria within one near-infrared cyanine, CyC4, encode two mechanistically distinct ratiometric reporting channels. A meso-amino group and a pendant carboxylate form a tunable intramolecular hydrogen bond that toggles the dye between closed (700 nm) and open (780 nm) emissive conformers. Time-dependent density functional theory (TD-DFT) calculations show that the hydrogen bond raises the LUMO and blue-shifts the emission, establishing the 700/780 emission ratio as a local reporter of hydrogen bonding and polarity. Independently, the chromophore self-associates under crowding- and cosolvent-rich conditions into an aggregate with a blue-shifted, H-type absorption signature near 530–540 nm and a distinct emission near 610 nm upon 540 nm excitation. The intensity of this aggregate band relative to the monomer emission (*R*_a_) serves as a ratiometric reporter of crowding and self-association. Because the two channels arise from distinct molecular equilibria (intramolecular hydrogen bonding vs. intermolecular self-association) they are largely decoupled: a glycerol titration series confirms that the self-association channel (*R*_a_) can be moved while the hydrogen-bonding channel stays essentially fixed. Applied to protein–PEG biomolecular condensates, the two ratios move oppositely with increasing salt, showing that the interior’s chemical (polarity, hydrogen bonding) and physical (packing, self-association) environments co-vary across the salt series; a single CyC4 measurement thereby maps this coupled microenvironment, providing a general strategy for multiparametric, ratiometric sensing of crowded microenvironments.

TOC Graphic

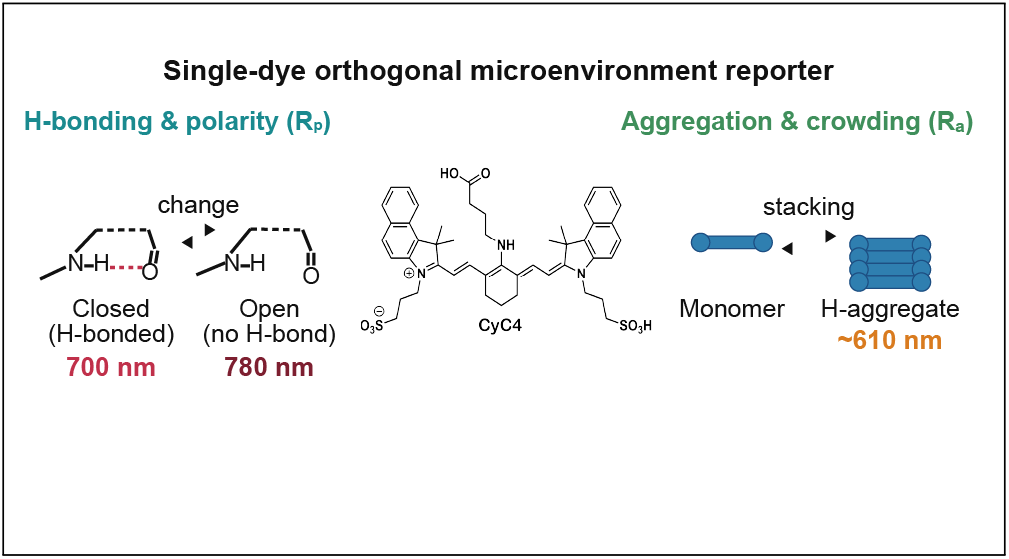

## Introduction

Environment-sensitive fluorophores report on their surroundings by convolving polarity, hydrogen bonding, viscosity, and self-association into a single spectral response.^1,2^ This degeneracy limits what a fluorescence change means in a single measurement: an emission shift recorded inside a cell or an organelle could reflect lower polarity, stronger hydrogen bonding, tighter molecular packing, or aggregation, making interpretation difficult. Reading more than one property from a single probe requires a chromophore in which distinct microenvironmental variables act through distinct, structurally defined channels that can be measured independently—ideally ratiometrically, and in the near-infrared (NIR) window where biological media scatter and absorb least.^3,4^

Heptamethine cyanines are attractive starting points for such a design. They possess high molar absorptivities and excellent brightness in the NIR region. Furthermore, their chemical versatility—allowing orthogonal functionalization at the indolenine nitrogen atoms and the central *meso* position of the polymethine chain—has established them as the preeminent class of NIR fluorescent probes.^5,6^ Their photophysical properties can respond to the surrounding microenvironment via multiple structurally and mechanistically distinct pathways. For instance, intramolecular charge transfer (ICT) and local conformational twisting modulate monomer emission profiles,^7,8^ while excitonic coupling during self-assembly into H- or J-type aggregates induces pronounced blue- or red-shifts in absorption and emission, respectively. ^9^ Critically, the former pathway represents an intramolecular reporter that alters the photo-physics of a single chromophore through charge redistribution and conformational gating, whereas the latter represents an intermolecular reporter driven by supramolecular interactions. Despite this dual capability, environment-sensitive cyanine probes have traditionally exploited these mechanisms in isolation—acting strictly as molecular rotors, aggregation sensors, or encapsulation-activated labels^1^—rather than integrating both mechanistically distinct modalities into a single molecular entity.

Here we show that a single meso-amino heptamethine cyanine, CyC4, achieves this dual-sensing objective by exploiting coupled chemical and physical molecular equilibria. An intramolecular hydrogen bond between the meso-amine and a pendant carboxylate interconverts two distinct emissive conformers (700 and 780 nm); TD-DFT calculations attribute the 700 nm band to hydrogen-bond-induced destabilization of the LUMO, and a chain-length synthetic control fixes the hydrogen bond as the switch, allowing the 700/780 emission ratio to report on hydrogen bonding and polarity as a ratiometric channel. Independently, the chromophore self-associates into an aggregate with a blue-shifted, H-type absorption signature under crowding- and cosolvent-rich conditions, giving rise to a distinct 610 nm emission upon 540 nm excitation, whose intensity relative to the monomer emission reports self-association and crowding. Because the two channels are governed by different molecular equilibria—one intramolecular, one intermolecular—they constitute two mechanistically distinct reporters that can be read out from a single probe through sequential, spectrally resolved excitation– emission measurements. Applied to protein–PEG phase-separated condensates^10–12^ whose interior polarity, packing, and hydrogen-bonding character are otherwise difficult to measure simultaneously in situ^13,14^—CyC4 resolves salt-dependent gradients within individual droplets and shows that the interior chemical and physical environments co-vary. Coupled molecular equilibria in a single chromophore thus provide a general strategy for multiparametric, ratiometric NIR sensing of crowded microenvironments.

## Results and discussion

### Synthesis and ground-state photophysics of CyC4

CyC4 is a cyclohexenyl-bridged, meso-amino heptamethine cyanine constructed on the bis-benzoindolium tricarbocyanine (Cy7) scaffold (Figure 1a). The defining structural feature is a 4-aminobutanoic acid side chain appended at the cyclohexenyl meso position: the four-carbon tether places the terminal carboxylate precisely within reach of the meso-amine N–H, enabling formation of a seven-membered intramolecular hydrogen-bonded ring. The two indolinium nitrogens carry propane-1-sulfonate groups that confer aqueous solubility without perturbing the polymethine chromophore. As a structural control, CyC6 was synthesized on the same scaffold with a six-carbon (6-aminohexanoic acid) tether; the additional two methylene units prevent closure of an intramolecular hydrogen bond, providing a direct comparison species that differs from CyC4 solely in H-bonding capacity.

**Figure 1:**
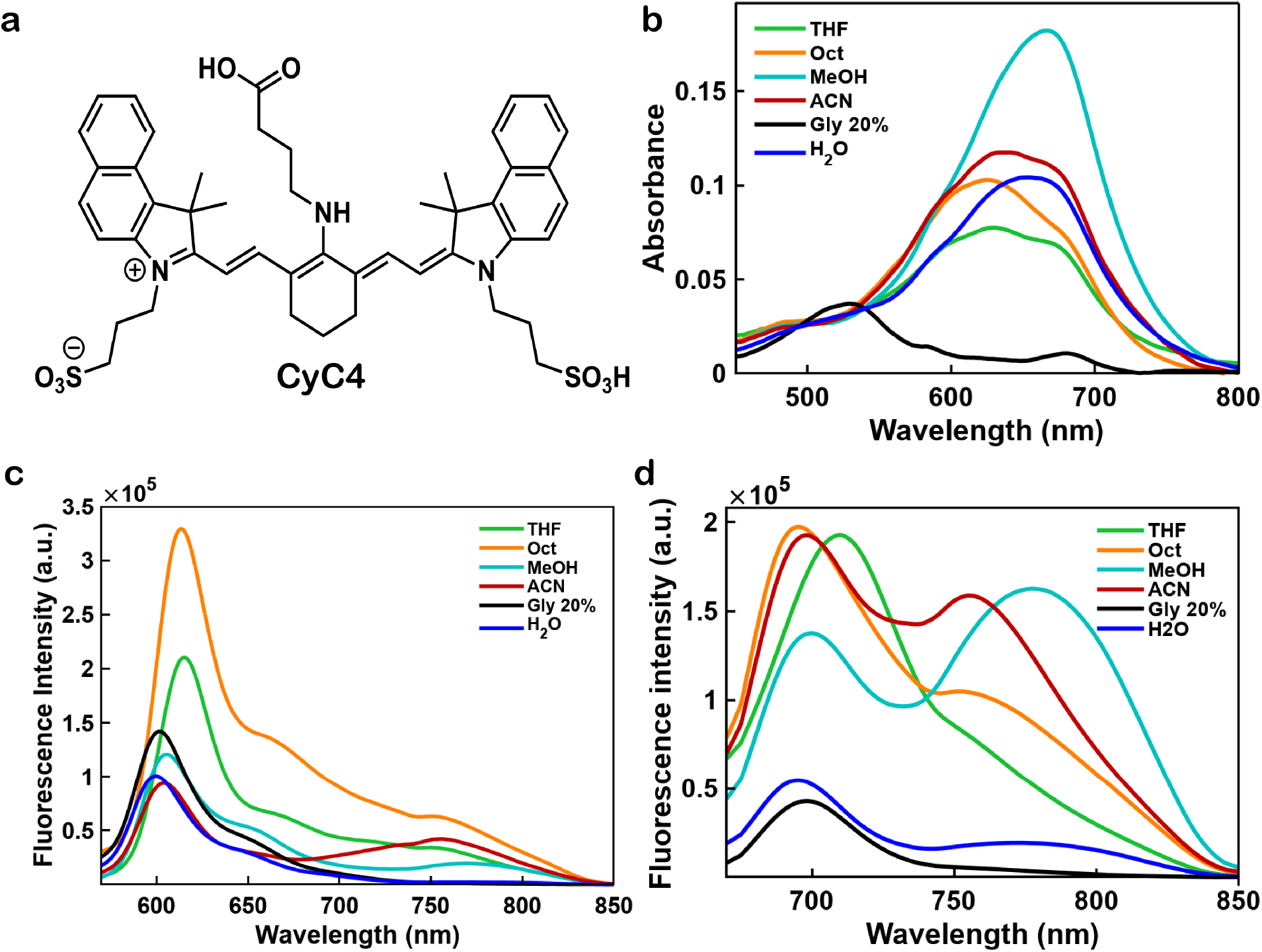
Structure and photophysics of CyC4. (a) Structure of CyC4: a cyclohexenyl-bridged, meso-amino heptamethine (Cy7) cyanine bearing a four-carbon carboxylate tether at the meso amine and propane-1-sulfonate groups on the indolenine nitrogens (the six-carbon control CyC6 differs only in tether length; see Scheme S1). (b) Absorption spectra of CyC4 (2.53 µM) across solvents of varying polarity and hydrogen-bonding character (tetrahydrofuran (THF), 1-octanol, methanol, acetonitrile, 20% glycerol, water); in 20% glycerol a strongly blue-shifted band near 530 nm appears, the ground-state signature of the self-associated aggregate. (c) Emission of CyC4 (0.569 µM) excited at 540 nm (*×*10^5^), isolating the blue-shifted (*∼*610 nm) aggregate-derived emission. (d) Emission of CyC4 (0.569 µM) excited at 660 nm (*×*10^5^), showing the 700 and 780 nm monomer bands (C-/O-states) whose relative intensity varies with solvent.

Both dyes were prepared in a single step from the meso-chloro precursor CyCl and the appropriate amino acid, and characterized by ^1^H NMR and high-resolution ESI-MS (Figures S1–S4; full synthetic and characterization details in the Experimental section and Supporting Information).

Ground-state molar extinction coefficients were determined in methanol by linear regression of the Beer–Lambert plot over 0.899–14.4 µM for CyC4 and 1.10–17.7 µM for CyC6 (Figure S5): *ε*_660_ = (4.30 *±* 0.02) *×* 10^4^ M^-1^cm^-1^ (SE, *n* = 14) for CyC4 and *ε*_673_ = (5.79 *±* 0.02) *×* 10^4^ M^-1^cm^-1^ (SE, *n* = 11) for CyC6. The difference in *ε* between CyC4 and CyC6, despite an identical chromophore, likely reflects differences in the ground-state conformational ensemble (vide infra): the H-bonded C-state of CyC4 places the carboxylate near the amine and distorts the planarity of the meso bridge relative to the predominantly open conformational ensemble of CyC6, reducing the transition dipole moment. Fluorescence quantum yields referenced to 3,3*^′^*-diethyloxatricarbocyanine iodide (DOTCI, Φ_F_ = 0.28 in methanol)^15^ were Φ_F_ = 0.041 for CyC4 and 0.039 for CyC6, demonstrating that neither the chain length nor the intramolecular H-bond materially reduces the radiative efficiency of the chromophore.

The absorption maximum of CyC4 shifts to the red with increasing solvent polarity (Figure 1b), a positive solvatochromism opposite to the classical negative solvatochromism of simple cyanines.^16^ This is consistent with a ground-state dipole moment that is *smaller* than that of the Franck–Condon excited state, so that polar solvents stabilize S_1_ relative to S_0_ and reduce the excitation energy.^17^

That the absorption energy is not governed solely by bulk polarity is evident from absorption spectra of CyC4 in acetonitrile and methanol: they have comparable permittivity (*ε_r_ ≈* 37.5 and 32.7) yet absorption maxima 26 nm apart (640 vs. 666 nm). This difference reflects specific hydrogen bonding in the protic solvent, demonstrating pronounced sensitivity of the absorption energy to both bulk polarity and specific solute–solvent hydrogen bonding. Additionally, in 20% glycerol, CyC4 develops a strongly blue-shifted band near 530 nm (Figure 1b), the ground-state signature of the self-associated aggregate characterised below.

### Dual NIR emission of CyC4 and the C/O two-state model

When CyC4 (0.569 µM) was excited at 660 nm, steady-state emission spectra revealed behavior unprecedented for heptamethine cyanines: two well-resolved emission bands at approximately 700 and 780 nm whose *relative* intensities vary strongly with the solvent environment (Figure 1d). The 700 nm band is most intense in THF and remains dominant in water and octanol; in acetonitrile it modestly exceeds the 780 nm band; in methanol the 780 nm band predominates (Figure 2c). The energy spacing between the two maxima (700 vs. 780 nm) is *≈*1470 cm*^−^*^1^, comparable to a conjugated C–C stretching frequency (intermediate between C–C single- and C=C double-bond stretches) and thus within the range of a 0–0/0–1 vibronic progression.^18^ The band separation alone therefore cannot establish whether the two bands are vibronic satellites of a single emitter or two distinct emissive states; the strong, solvent-dependent redistribution of intensity between them points to the latter, which the following experiments confirm.

**Figure 2:**
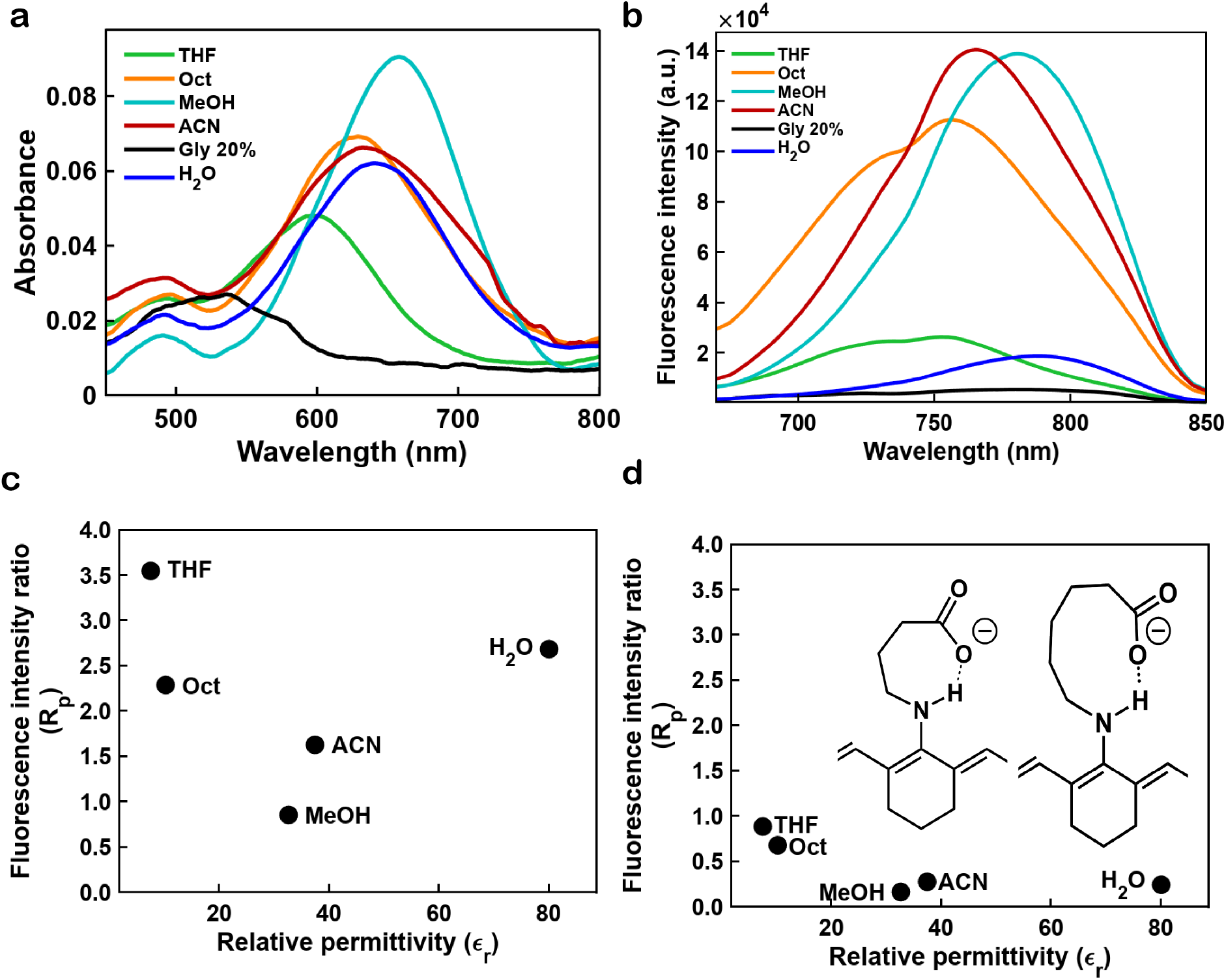
The polarity/hydrogen-bonding ratiometric channel of CyC4. (a) Absorption and (b, *×*10^4^) 660 nm-excited emission of CyC6 (1.05 µM and 0.246 µM, respectively) across solvents (tetrahydrofuran (THF), 1-octanol, methanol, acetonitrile, 20% glycerol, water). Unlike CyC4, CyC6 shows no resolved 700 nm band; emission is dominated by the 780 nm (O-state) band across all solvents. (c) The polarity/hydrogen-bonding ratio *R*_p_ = *I*_700_*/I*_780_ for CyC4, plotted against solvent relative permittivity *ε*_r_; *R*_p_ does not track permittivity monotonically, consistent with a hydrogen-bonding rather than a bulk-polarity origin. (d) The same *R*_p_–permittivity trend for CyC6, annotated with the DFT intramolecular hydrogen-bonded macrocyclic conformers of CyC4 and CyC6 (carboxylate O*· · ·* H–N), which rationalize the closed C-state.

Three independent experiments rule out the main alternative explanations. *First*, to distinguish two states from vibronic structure, excitation spectra were recorded while monitoring emission separately at 700 and 780 nm (Figure S6a,b). The excitation spectrum measured at *λ*_em_ = 780 nm closely mirrors the absorption spectrum, as expected for a single dominant species. In contrast, the excitation spectrum at *λ*_em_ = 700 nm revealed a distinct profile with additional features at *∼* 630 and 580 nm not present in the absorption spectrum. Because vibronic structure must give *identical* excitation spectra regardless of the monitored emission wavelength, these non-mirror-image excitation profiles exclude vibronic coupling as the origin of the dual-band emission and establish that the two bands arise from spectroscopically distinct ground-state populations. *Second*, concentration-dependent emission spectra recorded over a 14-fold range (0.0357–0.569 µM) in methanol, acetonitrile, water, and octanol yielded a monotonic increase in emission intensity with concentration, without the appearance of new emission bands, in all solvents (Figure S7). Within this dilute, single-solvent regime the 700/780 band ratio is independent of concentration, so the *monomer* C/O equilibrium is not driven by self-association. This does not exclude aggregation under other conditions: as shown in the next section, a distinct self-associated aggregate of CyC4 forms when crowding, cosolvent, or reduced temperature promote self-association, and that species is treated separately as the basis of the second (crowding) sensing channel. The structural origin of the dual emission is revealed by comparison with the chain-length control CyC6 (Figure 2a,b and Figure S6c). Under identical conditions (*λ*_ex_ = 660 nm), CyC6 produces a *single* emission maximum at *∼*780 nm in all solvents; its excitation spectrum at 780 nm mirrors the absorption with no additional features. Because CyC4 and CyC6 differ only in whether the carboxylate chain can geometrically close an intramolecular hydrogen-bonded ring, this comparison implicates the intramolecular H-bond as the structural switch responsible for the 700 nm band. The four-carbon linker of CyC4 is the minimum length required to form a seven-membered ring; the six-carbon linker of CyC6 cannot close the same seven-membered ring and strongly disfavors the compact, H-bonded C-state, although longer or solvent-bridged contacts are not excluded (SI). We therefore assign the 700 nm emission to the **C-state** (closed, H-bonded conformer) and the 780 nm emission to the **O-state** (open, non-H-bonded conformer). The emission ratio *I*_700_*/I*_780_ serves as an empirical ratiometric proxy for the C/O emission balance as a function of local environment; its calibration against solvent permittivity defines the polarity channel *R*_p_ (Figure 2c); the corresponding *R*_p_–permittivity trend for the CyC6 control, annotated with the DFT closed-conformer geometries for both dyes, is shown in Figure 2d.

Normalized emission spectra of CyC4 and CyC6 in each solvent confirm that the peak positions within each band are essentially solvent-invariant (*<*5 nm variation), while only the relative band amplitudes change with solvent (Figure S8). This rules out continuous solvatochromic shifts of a single state and is fully consistent with a two-state model in which each emitter has a fixed frequency and only the population balance changes. The non-monotonic relationship between *I*_700_*/I*_780_ and solvent polarity — highest in THF, high in water and octanol, and lowest in methanol (Figure 2c) — suggests that the C/O equilibrium is governed not by polarity alone but by the interplay of H-bond enthalpy, conformational entropy, and specific solute–solvent interactions, a picture elaborated by the computational analysis below.

### Computational basis for the two-state emission mechanism

To establish an atomic-level description of the C/O equilibrium and explain the counter-intuitive solvent dependence of *I*_700_*/I*_780_, we combined conformational ensemble generation with DFT and TD-DFT calculations in implicit solvent models (Figure 3). Full computational details are in the Supporting Information.

**Figure 3:**
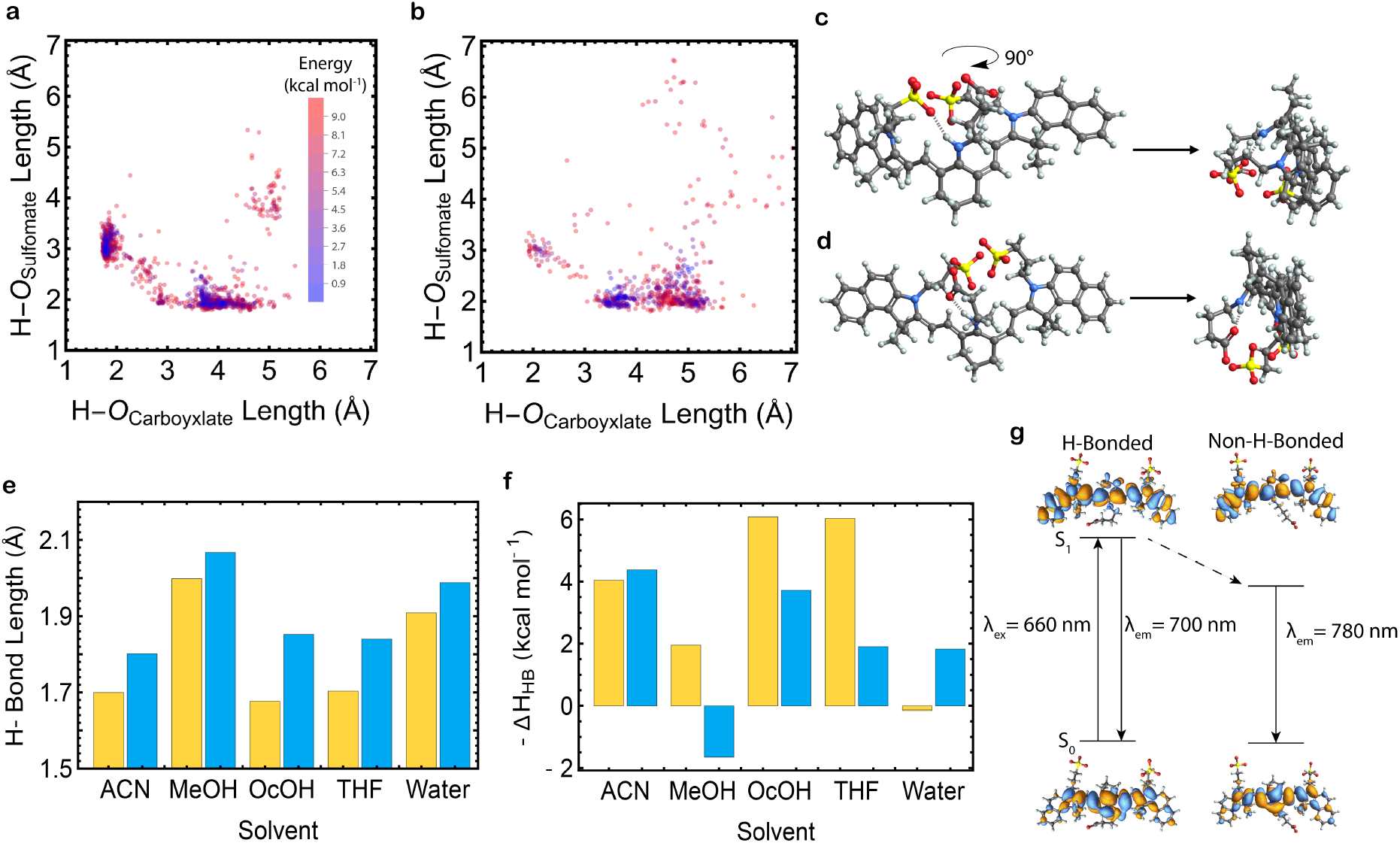
Computational basis of the intramolecular hydrogen-bond switch. (a,b) GOAT-generated CyC4 (a) and CyC6 (b) conformational ensembles in methanol plotted by intramolecular H*· · ·* O_carboxylate_ versus H*· · ·* O_sulfonate_ distances and colored by relative energy. (c,d) Lowest-energy members of the closed (carboxylate-H-bonded, C-state; c) and open (O-state; d) families of the CyC4-in-methanol ensemble, each shown in two orthogonal views (90*^◦^* rotation). (e) Intramolecular H-bond lengths per solvent for CyC4 (yellow) and CyC6 (blue) calculated by DFT at the *ω*B97X-D4/def2-TZVP level. (f) Corresponding H-bond enthalpies (*−*Δ*H*_HB_) per solvent, showing that solvent-dependent stabilization sets the C/O branching. (g) Jablonski scheme of the H-bonded (700 nm, C-state) and non-H-bonded (780 nm, O-state) emissive channels, with the corresponding HOMO and LUMO wavefunction isosurfaces.

Applying the Global Optimization Algorithm (GOAT) method^19^ in entropy maximization mode and using the GFN2-xTB Hamiltonian for CyC4 in methanol yields two Boltzmann-weighted conformer families—a short-H-bonded C-state and a broader open O-state (Fig ure 3a,b)—whereas CyC6 gives a single open distribution with no low-energy H-bonded cluster; representative optimized geometries are shown in Figure 3c,d, and the full set of DFT-optimized geometries in methanol is provided in Figure S9. DFT optimization (*ω*B97X-D4/def2-TZVP, ORCA; CPCM; DRACO when available) confirms that CyC4 forms shorter intramolecular hydrogen bonds than CyC6 in all three solvents (Figure 3e): H*· · ·* O_carboxylate_ distances in the CyC4 C-state are 1.82, 1.87, and 1.67 Å (water, methanol, acetonitrile) versus 2.05, 2.15, and 1.75 Å for CyC6. The ground-state intramolecular hydrogen bond is enthalpically favorable in acetonitrile (*−*Δ*H*_H-bond_ *≈* 4 kcal mol^-1^ for both dyes), weaker in methanol, and essentially neutral in water (*≈* 0 kcal mol^-1^ for CyC4 and 1.8 kcal mol^-1^ for CyC6), where competing solvent hydrogen bonds offset it; in every case it becomes less favorable in S_1_ (by up to *≈*5 kcal mol^-1^; Figure 3f, Table S3). This excited-state weakening sets the branching to the C-state (700 nm) or O-state (780 nm) minima (Figure 3g).

The 80 nm (*≈*0.18 eV) C/O emission gap follows from the orbital analysis. The HOMO is delocalized over the polymethine chain with negligible density at the meso amine (*−*6.94 eV in water, essentially H-bond-independent), whereas the LUMO has significant amplitude at the meso nitrogen. H-bond formation brings the anionic carboxylate near this electron-rich region and destabilizes the LUMO by *≈*0.15 eV (*−*0.78 *→ −*0.64 eV in water), widening the HOMO–LUMO gap and blue-shifting the C-state to 700 nm without any change in chromophore connectivity; TD-DFT vertical excitations reproduce the trend (H-bonded CyC4 451 nm vs. non-H-bonded 472 nm in water; complete TD-DFT excitations and frontier-orbital energies for both dyes across solvents in Table S4). The calculations are not intended to reproduce the quantitative *R*_p_ ordering across the full solvent series—which convolves the ground-state C/O population with the differential absorption and radiative yields of the two conformers and with specific solvent hydrogen bonding—but they capture its physical origin: the intramolecular hydrogen bond is preserved in aprotic media such as THF and progressively disrupted by competing solvent hydrogen bonds in protic media, while the excited state further favors the extended O-state.

### A third, aggregation-derived state and the crowding channel

The C/O two-state model accounts for the 660 nm-excited monomer emission but is incomplete. Exciting CyC4 at 540 nm—into the blue-shifted band that is weak in polar solvents yet dominant in glycerol-rich media (Figure 1b)—produces an emission envelope near 610 nm distinct from both monomer bands (Figure 1c), and the emission line shape depends on excitation wavelength (Figure S10). This excitation-wavelength-dependent emission—a signature of ground-state heterogeneity rather than of a single emitter—requires at least one further ground-state population beyond the C- and O-states.

This species is a self-associated aggregate with a blue-shifted, H-type absorption signature, not a solvatochromically shifted monomer: its absorption is blue-shifted (Figure 1b) and its population grows with glycerol and other cosolvents (Figure S10) and, more slowly, on standing (Figure S11)—hallmarks of self-association driven by reduced solvent quality, viscosity, and molecular crowding rather than by dye concentration, reconciling with the concentration invariance noted above. The blue-shifted absorption is consistent with H-type excitonic coupling, and the distinct 540 nm-excited emission indicates that this self-associated population is spectroscopically distinct from the monomeric C and O states.^20–22^ Crucially, the aggregate is a property of the chromophore, not the H-bond: CyC6—which cannot form the intramolecular H-bond—develops the same blue-shifted absorption and 540 nm-excited emission in glycerol-rich media, and the aggregate emission position is independent of protic/aprotic character. The aggregate axis is therefore largely decoupled from the hydrogen-bonding axis, as indicated by the *R*_a_-versus-permittivity calibration (Figure S10a).

We accordingly define two ratiometric observables: the hydrogen-bonding/polarity channel *R*_p_ = *I*_700_*/I*_780_ (Figure 2c) and the crowding/self-association channel

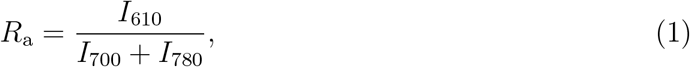

the 540 nm-excited aggregate emission (*I*_agg_) referenced to the *total* monomer emission. Referencing to the summed monomer population makes *R*_a_ insensitive to the C/O partition, so that the C/O partition is largely suppressed on the crowding axis. Normalization alone does not prove full independence—*R*_a_ still contains the monomer intensities in its denominator— but a glycerol titration provides direct evidence that the two respond separably (below). A glycerol titration shows that the aggregate emission *I*_610_ follows a Förster–Hoffmann relationship with viscosity (*R*^2^ = 0.90; viscosities from ref;^23^ Figure S10d), and *R*_a_ rises correspondingly with viscosity (Figure S10f), over a glycerol/water series in which viscosity co-varies with composition, establishing *R*_a_ as a reporter of self-association that responds to local crowding and solvent quality (a net balance) rather than to viscosity alone.

### Ratiometric sensing of biomolecular condensate microenvironments

Having established the two-state H-bonding mechanism and its molecular basis, we evaluated CyC4 as a ratiometric sensor of biomolecular condensate interiors. As a well-defined model system with experimentally controllable interior composition, we chose bovine serum albumin (BSA) condensates formed by liquid–liquid phase separation (LLPS) in the presence of PEG-8k and NaCl in sodium phosphate buffer. ^24,25^ In this system, NaCl concentration controls protein–protein electrostatic screening: as [NaCl] increases, BSA packing density in the dense phase increases, the water content decreases, and the interior microenvironment shifts from a dilute, water-like polarity toward a denser, intermediate-polarity milieu.^14,24^

BSA condensates were prepared at five NaCl concentrations (10, 50, 100, 150, and 200 mM) with fixed BSA (200 µM), PEG-8k (30 wt%), and sodium phosphate (100 mM, pH 7.0 ± 0.2), and incubated with 3.16 µM CyC4. CyC4 partitioned selectively and strongly into the dense phase, producing condensate fluorescence that exceeded the dilute-phase background by more than tenfold, indicating preferential uptake and enrichment.

Spectral images were acquired on a confocal microscope: the monomer channels by 660 nm excitation with simultaneous detection in 40 nm windows centred at 700 nm (C-state) and 760 nm (O-state), and the aggregate channel by a sequential 540 nm excitation, with emission collected across 550–790 nm (12 sequential 20 nm slices) and the 590–630 nm window (slices 2–3, averaged) used as the aggregate emission signal. Because the two excitation channels were acquired sequentially from the same field, the *R*_p_ and *R*_a_ images were spatially registered. Ratios were computed on background-subtracted intensities after per-image segmentation of the condensates and are reported as per-condensate means (full image-analysis pipeline in the Supporting Information): the polarity/hydrogen-bonding ratio *R*_p_ = *I*_700_*/I*_760_ and the aggregation/self-association ratio *R*_a_ = *I*_610_*/*(*I*_700_ + *I*_760_) (Eq. 1).

Pixel-wise *R*_p_ and *R*_a_ maps for representative condensates across the NaCl titration are shown in Figure 4a alongside the background-subtracted *I*_700_, *I*_760_, and *I*_610_ channels. Individual condensates display spatial heterogeneity in the ratio maps, with interior pixels differing from peripheral pixels in representative sections; we present these as qualitative maps and do not quantify radial profiles here.

**Figure 4:**
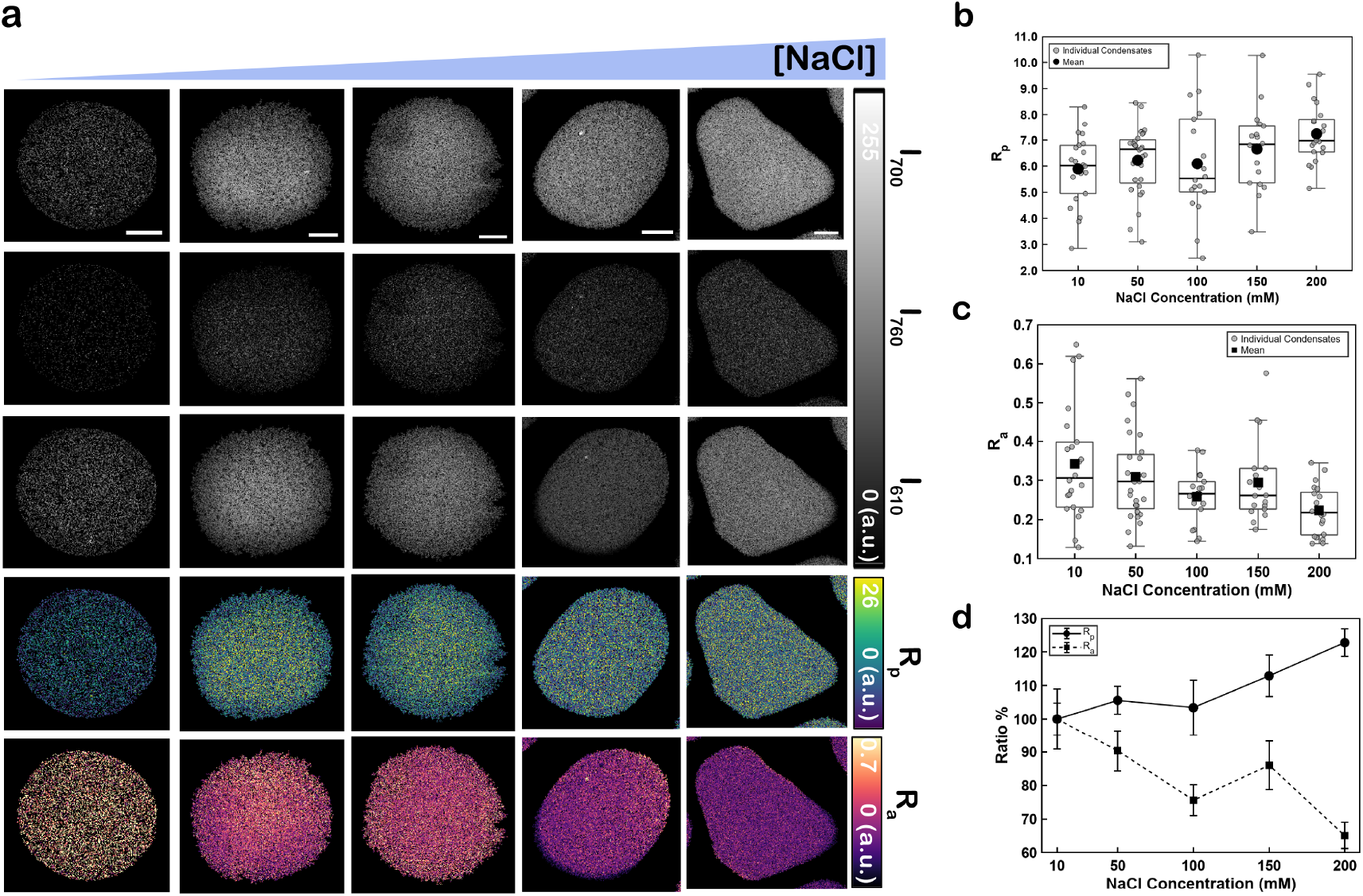
Dual-channel ratiometric mapping of BSA–PEG condensate microenvironments with CyC4. (a) [NaCl] titration (10, 50, 100, 150, 200 mM) of BSA condensates (200 µM BSA, 30 wt% PEG-8k, 100 mM sodium phosphate, pH 7.0 ± 0.2; 3.16 µM CyC4). For each condition the background-subtracted raw channels *I*_700_, *I*_760_ (660 nm excitation) and *I*_610_ (540 nm excitation) are shown together with the pixel-wise polarity-ratio map *R*_p_ = *I*_700_*/I*_760_ and aggregation-ratio map *R*_a_ = *I*_610_*/*(*I*_700_ + *I*_760_). Scale bar 10 µm. (b) Per-condensate *R*_p_ versus [NaCl] (box-and-whisker over individual condensates, grey; mean, black). (c) Per-condensate *R*_a_ versus [NaCl] (same convention). (d) Mean *R*_p_ and *R*_a_ normalized to their 10 mM values (%), showing that *R*_p_ increases and *R*_a_ decreases with ionic strength. *n* = 22, 28, 18, 18, 21 condensates at 10, 50, 100, 150, 200 mM NaCl across three biological replicates. Error bars represent SEM across individual condensates within each condition. The two ratios move in opposite, consistent directions across the full NaCl range (Spearman rank correlation with [NaCl] on the per-condition means in panel d: *R*_p_ *ρ* = +0.90; *R*_a_ *ρ* = *−*0.90).

Per-condensate mean *R*_p_ and *R*_a_ across the salt series are quantified in Figure 4b and c, respectively, with each condensate plotted individually. The two channels moved in opposite directions across the salt series. The polarity ratio increased with ionic strength, from *R*_p_ = 5.90 *±* 0.28 at 10 mM NaCl to 7.24 *±* 0.24 at 200 mM, with intermediate values of 6.23 *±* 0.24 (50 mM), 6.10 *±* 0.49 (100 mM), and 6.66 *±* 0.37 (150 mM) (mean *±* SEM across individual condensates; *n* = 22, 28, 18, 18, 21 condensates across three biological replicates; condensates within a preparation are not independent). Over the same series the aggregation/self-association ratio decreased, from *R*_a_ = 0.343 *±* 0.031 at 10 mM to 0.223 *±* 0.013 at 200 mM (0.310 *±* 0.021, 0.259 *±* 0.016, and 0.295 *±* 0.025 at 50, 100, and 150 mM). Both trends are directional rather than strictly monotonic, with the 150 mM point departing slightly from the overall progression.

The two channels are mechanistically distinct and can be moved separately. In a glycerol titration *R*_a_ rises with viscosity while *R*_p_ is essentially constant (Figure S10f), so the self-association and hydrogen-bonding channels can be moved separately. This separation is expected for a glycerol cosolvent: glycerol builds an intermolecular hydrogen-bonding network that closely mimics that of water,^26^ so the titration raises viscosity while leaving the interior hydrogen-bonding character—and hence *R*_p_—largely unchanged. In the condensate salt series, by contrast, the two ratios move in *opposite* directions and are showed opposing salt-dependent trends (Spearman *ρ* = +0.90 and *−*0.90 on the five per-condition means, reported as a descriptor of monotonicity rather than an inferential statistic; Figure 4d). Read against the Figure 2 and Figure S10 calibrations, the rising *R*_p_ reports a chemical change: as interior water—a competing hydrogen-bond partner—is depleted with increasing packing, the intramolecular carboxylate–amine hydrogen bond that defines the C-state is less disrupted and emission shifts toward 700 nm. The falling *R*_a_ reports the coupled change in self-association. Although denser protein packing would be expected to increase the aggregate signal—much like viscosity does in the glycerol series—the accompanying compositional shift inside the condensate (such as water depletion and change in local dielectric properties) likely alters monomer solvation and shifts the aggregate–monomer equilibrium. Under these conditions, this solvation effect outweighs the viscosity-driven tendency toward aggregation, resulting in a net decrease in *R*_a_. The two reporters thus remain mechanistically distinct, but their opposite motion shows that the interior chemical (polarity, hydrogen bonding) and physical (packing, self-association) environments are not independent—they co-vary as the dense phase densifies. A single CyC4 measurement therefore resolves both microenvironmental axes and their correlation, directly visualizing the physical–chemical coupling that we recently inferred, by an independent lifetime-based approach, in a companion study of condensate interiors.^14^

The ratiometric readout confers a critical practical advantage over single-channel intensity measurements in these systems. BSA condensates vary in size, dye partitioning efficiency, and axial position within the confocal detection volume; these factors introduce condensate-to-condensate intensity variation of 30–50% that would confound any single-channel polarity measurement. The *I*_700_*/I*_760_ ratio reduces common-mode variability arising from condensate size, dye loading, and axial position, providing a self-normalized readout that is primarily sensitive to changes in the local C/O emission balance.

CyC4 is a near-infrared dual-emission cyanine whose two emissive states are governed by a single, rationally tunable intramolecular hydrogen bond (the carboxylate–amine interaction, which is strongly disfavored in the CyC6 linker-length control). This mechanism, supported by quantum-chemical calculations, is distinct from previously reported dual-fluorescent aminocyanines^27^ in that the two states are toggled by a single, structurally defined intramolecular contact. Moreover, CyC4 is multiparametric: the same molecule reports two mechanistically distinct microenvironmental axes—hydrogen bonding/polarity through *R*_p_ and self-association/crowding through *R*_a_—turning the otherwise-confounding aggregation of NIR cyanines into a second, mechanistically distinct ratiometric readout of the condensate interior. The design principle — placing a short-chain amino acid at the meso position of a tricarbocyanine to exploit excited-state electrostatic destabilization of the H-bonded conformer — is modular and adaptable: amino acids of different chain length will tune the ring size and H-bond geometry; fluorinated or caged carboxylates could shift the C/O pK*_a_*; and the Cy7 scaffold can be extended to heptamethine-to-nonamethine variants for longer-wavelength operation in the NIR-II window. The approach thereby opens a design space for ratiometric probes of biomolecular condensate interiors, organelle microenvironments, and other crowded biological compartments where existing single-wavelength dyes have limited interpretive power.

## Experimental

### Materials

All reagents and solvents were obtained from commercial suppliers and used as received unless otherwise noted. Methanol (ACS grade, CAS 67-56-1) and tetrahydrofuran (HPLC grade, CAS 109-99-9) were purchased from Fisher Chemical (Fair Lawn, NJ). 1-Octanol (ACS grade, CAS 111-87-5) was purchased from Sigma-Aldrich (St. Louis, MO). Acetonitrile (HPLC grade, CAS 75-05-8) was purchased from Crystalgen, Inc. (Commack, NY). Glycerol (ACS grade, CAS 56-81-5) was purchased from Research Products International (RPI, Mt. Prospect, IL). Ultrapure water (18.2 MΩ·cm) was obtained from a Milli-Q IQ 7000 water purification system (MilliporeSigma, Burlington, MA) and used to prepare all aqueous solutions. Bovine serum albumin (BSA, CAS 9048-46-8) was purchased from Tocris Bioscience (Bristol, UK). Polyethylene Glycol 8000 (PEG, MW 8000, CAS 25322-68-3) was purchased from Fisher BioReagents (Fair Lawn, NJ). Sodium chloride (CAS 7647-14-5) was purchased from Fisher Chemical (Fair Lawn, NJ). Sodium phosphate dibasic (CAS 7558-79-4, Sigma–Aldrich, St. Louis, MO) and sodium phosphate monobasic monohydrate (CAS 10049-21-5, Fisher Chemical, Fair Lawn, NJ) were used to prepare sodium phosphate buffer, as described below.

### Synthesis of CyC4 and CyC6

CyC4 and CyC6 were prepared by aromatic nucleophilic substitution (S_RN_1) of the central chlorine of the bis(benzoindolium) tricarbocyanine (CyCl) precursor with 4-aminobutanoic acid (CyC4) or 6-aminohexanoic acid (CyC6) in anhydrous DMF at 80 °C for 16 h under argon with no added base.^28^ After size-exclusion chromatography followed by reversed-phase preparative TLC, CyC4 and CyC6 were isolated as deep blue solids in 45% and 53% yield, respectively. Identity and purity were confirmed by ^1^H NMR (Bruker Avance II-500, 500 MHz), and high-resolution ESI-MS (Bruker Daltonik microTOF-Q); full synthetic procedures and characterization data are provided in the Supporting Information.

### Steady-state absorption and fluorescence spectroscopy

UV–vis absorption spectra were recorded from 300–800 nm on a single beam VWR UV-1600PC Spectrophotometer and steady-state fluorescence spectra on a HORIBA QM400 fluorescence spectrophotometer, using 1 cm quartz cuvettes at ambient temperature unless stated otherwise. Fluorescence spectra were acquired with excitation and emission slit widths of 2.5 nm, an integration time of 1 s and a step size of 1 nm unless otherwise stated. Absorption spectra were recorded using 2.53 µM CyC4 and 1.05 µM CyC6, and emission and excitation spectra using 0.569 µM CyC4 and 0.246 µM CyC6, unless otherwise stated. Molar extinction coefficients were determined in methanol by linear regression of the Beer– Lambert plot over 0.90–14.4 µM for CyC4 and 1.10–17.7 µM for CyC6. Monomer emission was excited at 660 nm (670–850 nm emission range) and the aggregate channel at 540 nm (550–850 nm emission range); excitation spectra were recorded over 490–690 nm while monitoring emission at 700 nm, 490–760 nm while monitoring emission at 780 nm, and 400–600 nm while monitoring emission at 610 nm. Fluorescence quantum yields were referenced to 3,3*^′^*-diethyloxatricarbocyanine iodide (DOTCI, Φ_F_ = 0.28 in methanol)^15^ with refractive-index correction. The solvent series comprised tetrahydrofuran, 1-octanol, methanol, acetonitrile, 20% glycerol solution in water and water. All spectra were blank/solvent-subtracted.

### Cosolvent- and time-dependent aggregation

Aggregation was probed by glycerol titration in water (0–20 v/v%, 5 v/v% steps) using 0.57 µM CyC4, with 540 nm excitation to follow the blue-shifted aggregate emission and 660 nm excitation to follow the monomer bands. The time dependence of aggregation at fixed composition was recorded in water for CyC4 (1.5 µM) and CyC6 (0.41 µM), also monitored with 540 and 660 nm excitation.

### Computational methods

Computations were performed using ORCA 6.1.0. ^29^ Conformational ensembles were generated using GOAT^19^ with the GFN2-xTB Hamiltonian and ddCOSMO solvation model in entropy mode with a convergence threshold of 0.05 cal mol*^−^*^1^ K*^−^*^1^, maximum energy of 10 kcal mol*^−^*^1^ with respect to the global minimum, and rotational constant difference of 0.1% to capture complete and diverse ensembles. ^30^ Geometries were optimized at the *ω*B97X-D4/def2-TZVP level^31^ with the CPCM^32^ implicit solvent model, using the DRACO radius scheme when available,^33^ in five solvents (tetrahydrofuran, 1-octanol, methanol, acetonitrile, and water). Harmonic frequencies were computed for water, methanol, and acetonitrile to obtain the ground- and excited-state hydrogen-bond thermochemistry (Table S3), the excited-state values estimated from the geometry-optimized ground states and TD-DFT^34^ vertical excitation energies at the same level; complete TD-DFT vertical excitations and frontier-orbital energies for all five solvents are given in Table S4. Conformer populations were Boltzmann-weighted at 298 K. Optimized Cartesian coordinates and total energies for all reported CyC4 and CyC6 conformers are provided as a separate Supporting Information archive (CyC4 CyC6 DFT optimized geometries.zip).

### BSA–PEG condensate preparation

100 mM sodium phosphate buffer (pH 7.0 ± 0.2) was prepared by mixing 39 mL of 1 M sodium phosphate dibasic (Na_2_HPO_4_) and 61 mL of 1 M sodium phosphate monobasic (NaH_2_PO_4_), followed by dilution to a final volume of 1.0 L with Milli-Q water. The pH was verified at room temperature and adjusted to 7.0 ± 0.2, if necessary, using dilute HCl or NaOH. Glass-bottom 384-well plates (Cellvis) were passivated with 1% Tween-20 for 1 h at 37 *^◦^*C, washed three times with Milli-Q water and three times with sodium phosphate buffer, and dried under a gentle stream of nitrogen prior to use. Condensates were formed at room temperature by liquid–liquid phase separation in 1.5 mL centrifuge tubes by sequential addition of sodium phosphate buffer, NaCl (10, 50, 100, 150, or 200 mM), BSA (200 µM final, from a 750 µM stock), CyC4 (3.16 µM final), and PEG-8k (30 wt% final, from a 50% stock). Condensates were transferred immediately to the passivated imaging plate and imaged 5 min after transfer, at room temperature.

### Confocal ratiometric imaging and analysis

Spectral images were acquired on a Leica TCS SP8 X confocal microscope equipped with a white-light laser, using a 100*×*, NA 1.44 oil-immersion objective and a 1-Airy-unit pinhole, at 512*×*512 pixels (0.227 µm/pixel). Condensates were excited at *λ*_ex_ = 660 nm (50% laser power), with simultaneous detection in two 40 nm windows centered at 700 nm (C-state) and 760 nm (O-state); the crowding channel uses an interleaved 540 nm excitation, with emission collected across 550–790 nm (12 sequential 20 nm slices), of which the 590–630 nm window (slices 2–3, averaged) was used as the aggregate emission signal for *R*_a_.

Condensates were segmented per image from the 700 nm channel via Otsu thresholding. Background was estimated per channel and subtracted from each condensate’s mean intensity post-segmentation. Ratios *R*_p_ = *I*_700_*/I*_760_ and *R*_a_ = *I*_610_*/*(*I*_700_ + *I*_760_) were computed from background-subtracted, per-condensate mean channel intensities, and reported as per-condensate values. Because the confocal detector and the solution fluorimeter have different spectral responses, the O-state emission is collected in a 760 nm window on the microscope versus at the 780 nm band maximum in solution; the two *R*_p_ definitions are therefore instrument-specific and not on the same absolute scale, so the imaging *R*_p_ is interpreted for its direction and relative change rather than its absolute magnitude. Condensates were analysed across three biological replicates (*n* = 22, 28, 18, 18, and 21 condensates for 10, 50, 100, 150, and 200 mM NaCl, respectively; each condition represented in all three replicates).

## Acknowledgement

The authors thank members of the Saurabh research group for helpful discussions. This study was supported by the NYU Discovery Research Funds to S.S., and by the Fondo para la Investigación Científica y Tecnológica (PICT 2021-252), the Consejo Nacional de Investigaciones Científicas y Técnicas (11220210100319CO), and the Secretaría de Ciencia y Técnica, Universidad de Buenos Aires (20020190100052BA) to C.S. This work was supported in part through the NYU IT High Performance Computing resources, services, and staff expertise.

## Supporting Information Available

Full synthetic procedures and ^1^H NMR and HR-ESI-MS characterization of CyC4, CyC6, and the synthetic intermediates; molar extinction-coefficient plots; solvent-dependent excitation and emission spectra; concentration-independence of the *I*_700_*/I*_780_ ratio; peak-normalized emission spectra; DFT-optimized geometries, intramolecular hydrogen-bond thermochemistry, and TD-DFT vertical excitations and frontier-orbital energies (Tables S3 and S4), with optimized Cartesian coordinates and energies provided as a separate archive; and the solvent-, viscosity-, and time-dependence of the aggregate/monomer balance.

