## Supplementary Information for "Multiparametric microenvironment sensing via distinct molecular equilibria in a single cyanine dye"

### Contents

**Synthesis and characterization** **S-3**

**Ground-state and steady-state photophysics** **S-11**

**Computational details** **S-15**

**Aggregation and the crowding channel** **S-17**

### Synthesis and characterization

Complete synthetic procedures and characterization for CyC4, CyC6, and the intermediates S1, S2, and CyCl are given below; the overall route is summarized in Scheme S1 and the intermediate syntheses in Scheme S2. All reagents and solvents were analytical grade (Sigma–Aldrich) and used as received unless otherwise stated. NMR spectra were recorded on a Bruker Avance II-500, 500 MHz spectrometer and mass spectra on a Bruker Daltonik microTOF-Q or a Waters Xevo G2-S Q-TOF instrument (ESI+/ESI−), as specified for each compound.

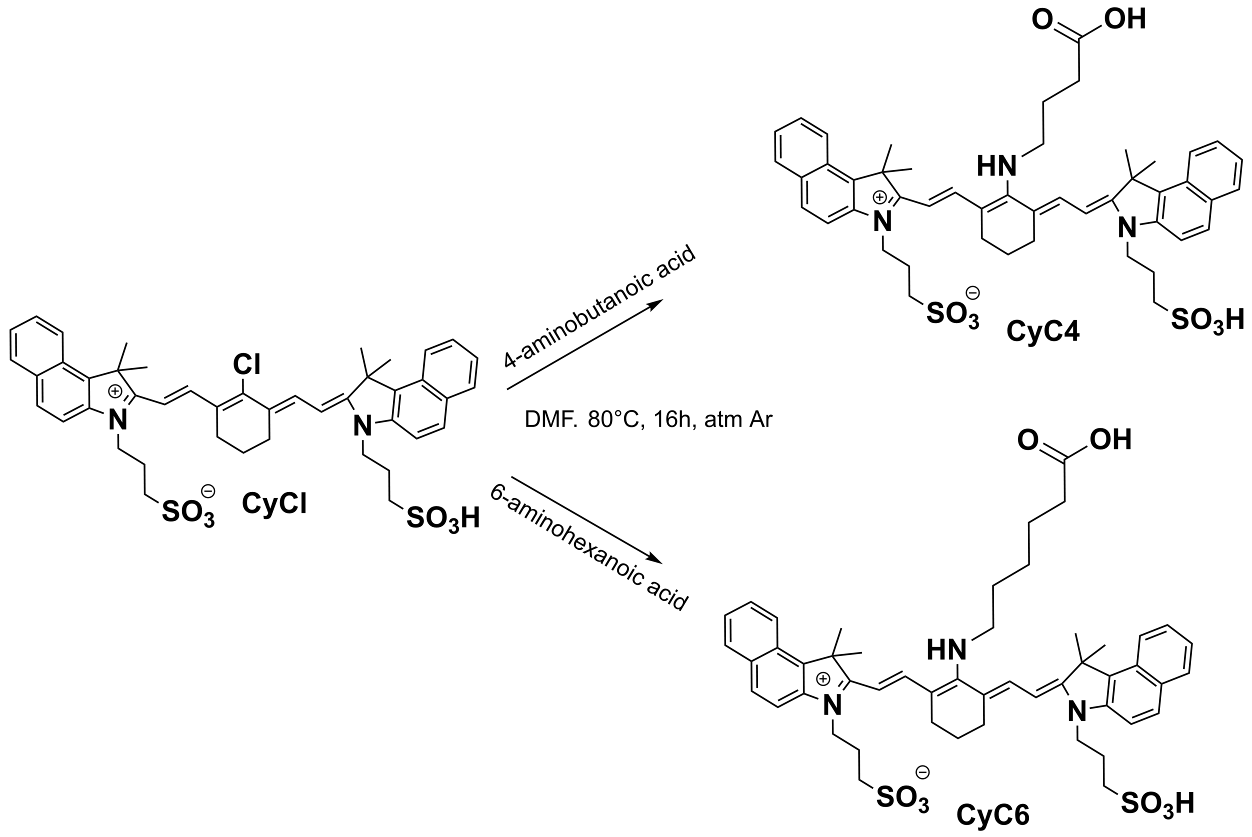


Scheme S1: Synthesis of CyC4 and CyC6 by aromatic nucleophilic substitution of the *meso*-chloro precursor CyCl with 4-aminobutanoic acid (CyC4) or 6-aminohexanoic acid (CyC6).

#### 1,1,2-Trimethyl-3-(3-sulfopropyl)-1H-Benzo[e]indolium-, inner Salt (S1)

Toluene (5 mL), 1,1,2-trimethylbenzo[e]indole (1.3 g, 6.2 mmol), and 1,3-propanesultone (1.1 mL, 12.5 mmol) were heated under reflux for 18 h. After cooling, the bluish crystals were filtered, washed with ether (3 × 10 mL), recrystallized from MeOH/$Et_2O$, and dried in vacuo (1.68 g, 82%). $^1$H NMR (200 MHz, MeOD-$d_4$) $\delta$ 8.30 (d, $J = 8.4$ Hz, 1H), 8.19 (d, $J = 9.0$ Hz, 1H), 8.11 (d, $J = 8.9$ Hz, 2H), 7.78 (t, $J = 7.1$ Hz, 1H), 7.68 (t, $J = 7.5$ Hz, 1H), 4.75 (t, $J = 7.2$ Hz, 2H), 3.30 (dd, $J = 3.2, 1.6$ Hz, 2H), 2.43 (dt, $J = 14.5, 7.7$ Hz, 2H), 1.82 (d, $J = 3.6$ Hz, 6H). HRMS (ESI+): calcd $(M+H)^+$ 332.1315; found 332.1315.

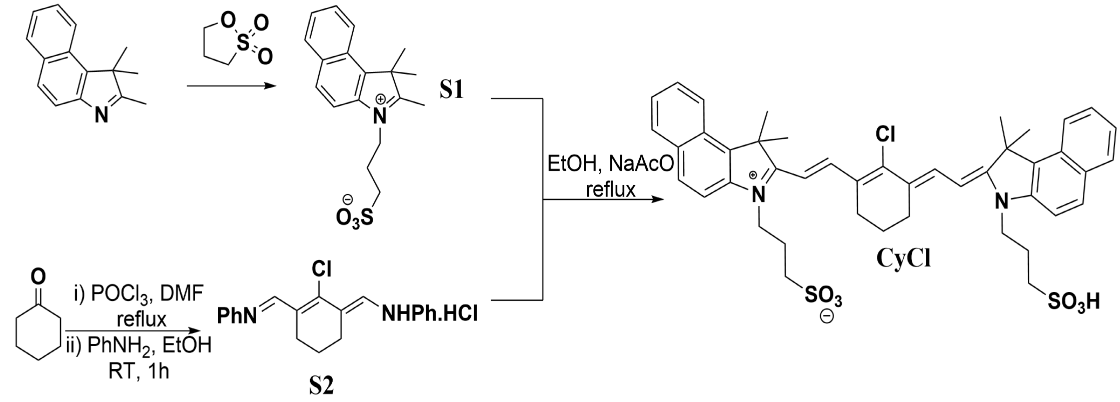


Scheme S2: Synthesis of the intermediates S1, S2, and CyCl.

#### N-[5-Anilino-3-chloro-2,4-(propane-1,3-diyl)-2,4-pentadiene-1-ylidene]anilinium Chloride (S2)

At 0 °C, phosphorus oxychloride (2 mL, 22 mmol) was added dropwise to anhydrous DMF (2.4 mL, 31 mmol). After 30 min, cyclohexanone (1 mL, 9.6 mmol) was added and the mixture refluxed for 1 h. An aniline/EtOH mixture [1:1 (v/v), 3.3 mL] was added dropwise at 20 °C; after a further 30 min the deep-purple mixture was poured into ice-cold $H_2O$/concentrated HCl (10:1, 20 mL). Crystals formed over 2 h in an ice bath were filtered, washed with cold $H_2O$ and $Et_2O$, and dried in vacuo (2.95 g, 84%). $^1$H NMR (200 MHz, MeOD-$d_4$) $\delta$ 8.68 (s, 2H), 7.58–7.17 (m, 10H), 2.74 (t, $J = 6.2$ Hz, 4H), 2.11–1.75 (m, 2H). HRMS (ESI+): calcd $(M+H)^+$ 323.1310; found 323.1308.

#### CyCl

A solution of S1 (1.69 g, 6 mmol), S2 (1.079 g, 3 mmol), and anhydrous sodium acetate (600 mg, 7 mmol) in absolute EtOH (60 mL) was heated under reflux for 3.5 h under $N_2$. The EtOH was removed under reduced pressure, the residue washed with $Et_2O$, and the crude recrystallized from MeOH/$Et_2O$ (1.35 g, 93%). $^1$H NMR (500 MHz, DMSO-$d_6$) $\delta$ 8.32 (d, $J = 14.1$ Hz, 2H), 8.25 (d, $J = 8.5$ Hz, 2H), 8.05 (d, $J = 8.9$ Hz, 2H), 8.01 (d, $J = 8.3$ Hz, 2H), 7.82 (d, $J = 9.0$ Hz, 2H), 7.60 (td, $J = 6.8, 1.2$ Hz, 2H), 7.47 (td, $J = 6.8, 1.2$ Hz, 2H), 6.52 (d, $J = 14.3$ Hz, 2H), 4.47 (t, 4H), 2.74 (t, $J = 5.6$ Hz, 4H), 2.60 (t, $J = 6.7$ Hz, 4H), 2.05 (quin, $J = 7.2$ Hz, 4H), 1.90 (s, 12H), 1.82 (quin, $J = 5.8$ Hz, 3H). HRMS (ESI−): calcd 797.2486; found 797.2493.

#### Synthesis of CyC4

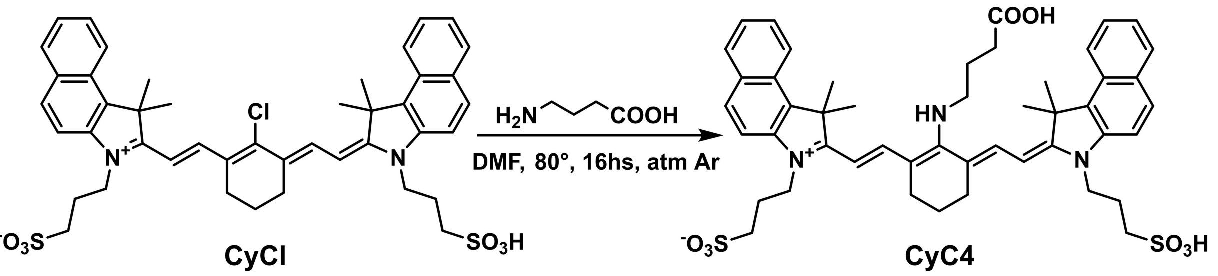


Scheme S3: Synthesis of CyC4 from CyCl and 4-aminobutanoic acid.

Following Scheme S3, CyCl (0.024 g, 0.03 mmol) and 4-aminobutanoic acid (0.0062 g, 0.06 mmol) were dissolved in anhydrous DMF (2 mL). The reaction mixture was stirred and heated at 80 °C for 16 h under argon atmosphere and protected from light. During the course of the reaction, the solution color gradually changed from green to deep blue, indicating successful dye formation. The solvent was removed under reduced pressure, and the crude product was purified by size-exclusion column chromatography (Sephadex LH-20, MeOH) followed by preparative thin-layer chromatography using a reversed-phase C18 stationary phase and methanol/water (7:3, v/v) as the mobile phase. The desired product was isolated as a deep blue solid in 45% yield (11 mg).

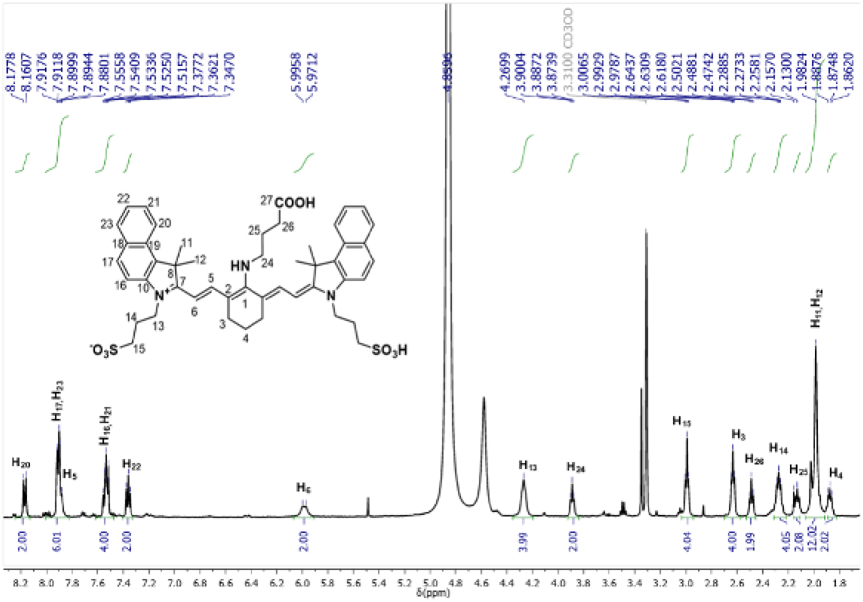


Figure S1: ${}^{1}$H NMR spectra of CyC4 (500 MHz, $CD_3OD$).

${}^{1}$H NMR (500 MHz, $CD_3OD$; Figure S1), $\delta$: 8.178 (d, 2H, $J$ = 8.5 Hz, $H_{20}$), 7.918 (dd, 4H, $J$ = 9.0, 3.0 Hz, $H_{17}$ and $H_{23}$), 7.880 (d, 2H, $J$ = 12.5 Hz, $H_5$), 7.556 (m, 4H, $H_{16}$ and $H_{21}$), 7.377 (t, 2H, $J$ = 7.5 Hz, $H_{22}$), 5.996 (d, 2H, $J$ = 12.5 Hz, $H_6$), 4.270 (m, 4H, $H_{13}$), 3.887 (t, 2H, $J$ = 6.5 Hz, $H_{24}$), 2.993 (t, 4H, $J$ = 6.5 Hz, $H_{15}$), 2.631 (t, 4H, $J$ = 6.5 Hz, $H_3$), 2.488 (t, 2H, $J$ = 7.0 Hz, $H_{26}$), 2.273 (t, 4H, $J$ = 7.5 Hz, $H_{14}$), 2.157 (m, 2H, $H_{25}$), 1.982 (s, 12H, $H_{11}$ and $H_{12}$), 1.875 (m, 2H, $H_4$).

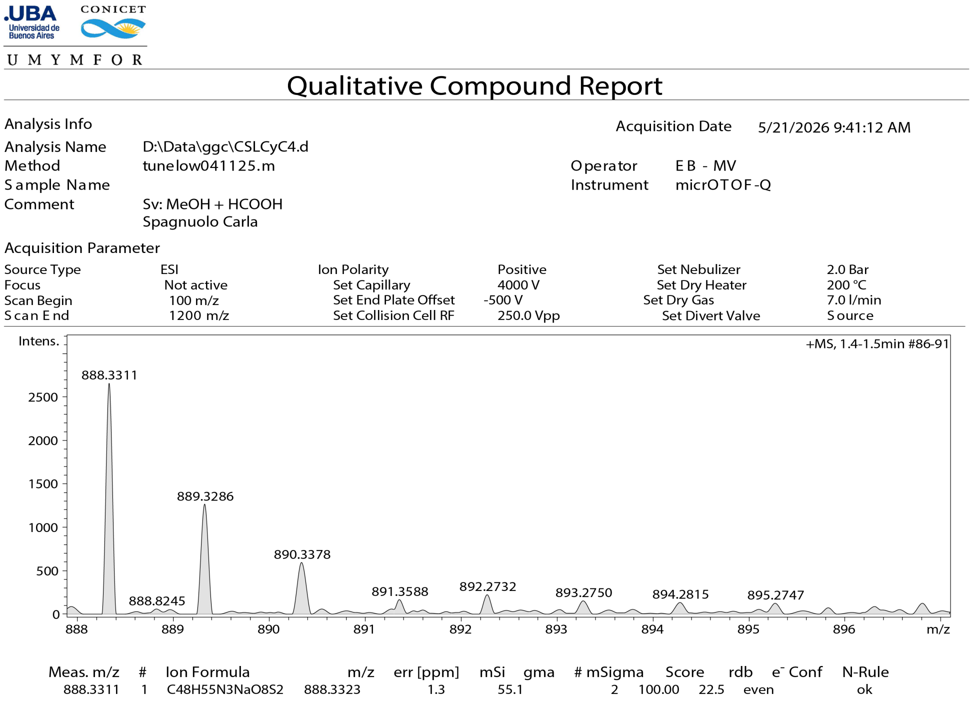


Figure S2: Representative high-resolution ESI mass spectrum of CyC4.

**HRMS (ESI$^+$)** calculated for $C_{48}H_{55}N_3NaO_8S_2$ $[M + Na]^+$: $m/z$ 888.3323; found: $m/z$ 888.3311 (Figure S2, Table S1).

Table S1: High-resolution ESI mass-spectrometry data for CyC4.

| $m/z$ experimental | Formula of ionic species | $m/z$ calculated | Proposed ionic species | Error (ppm) |
|---|---|---|---|---|
| 888.3311 | $C_{48}H_{55}N_3NaO_8S_2$ | 888.3323 | $[M + Na]^+$ | 1.3 |

LC–MS-grade reagents were used to prepare the sample solution. High-resolution mass spectrometry was performed by direct infusion using a microTOF-Q mass spectrometer equipped with an electrospray ionization (ESI) source. Data were acquired in positive-ion mode over an $m/z$ range of 100–1200. The $[M + Na]^+$ ion was detected at $m/z$ 888.3311, in agreement with the calculated value of 888.3323 (1.3 ppm mass error).

#### Synthesis of CyC6

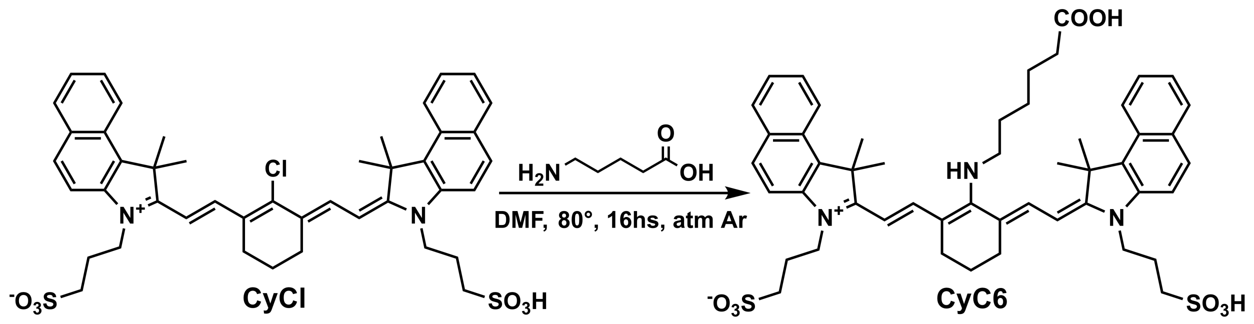


Scheme S4: Synthesis of CyC6 from CyCl and 6-aminohexanoic acid.

Following Scheme S4, CyCl (0.1 g, 0.13 mmol) and 6-amino hexanoic acid (0.033 g, 0.26 mmol) were dissolved in anhydrous DMF (5 mL) and the solution was heated at 80°C for 16 h protected from light and under argon atmosphere. The solution color gradually changed from green to deep blue. The solvent was removed under vacuum and the product was obtained as a deep blue solid in 53% yield (60 mg). The crude product was purified by size-exclusion column chromatography (Sephadex LH-20, MeOH) followed by preparative thin-layer chromatography using a reversed-phase C18 stationary phase and methanol/water (7:3, v/v) as the mobile phase.

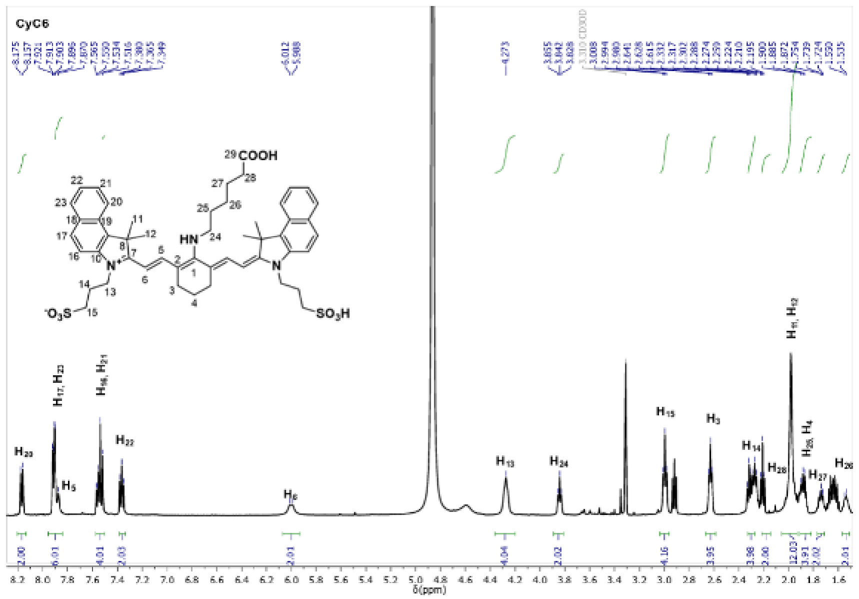


Figure S3: $^1H$ NMR spectra of CyC6 (500 MHz, $CD_3OD$).

$^1H$ NMR (500 MHz, $CD_3OD$; Figure S3), $\delta$: 8.175 (d, 2H, $J$ = 9.0 Hz, $H_{20}$), 7.921 (dd, 4H, $J$ = 9.0, 4.0 Hz, $H_{17}$ and $H_{23}$), 7.870 (d, 2H, $J$ = 12.5 Hz, $H_5$), 7.565 (m, 4H, $H_{16}$ and $H_{21}$), 7.380 (t, 2H, $J$ = 7.5 Hz, $H_{22}$), 6.012 (d, 2H, $J$ = 12.5 Hz, $H_6$), 4.273 (m, 4H, $H_{13}$), 3.855 (t, 2H, $J$ = 6.5 Hz, $H_{24}$), 3.008 (t, 4H, $J$ = 7.0 Hz, $H_{15}$), 2.641 (t, 4H, $J$ = 6.5 Hz, $H_3$), 2.302 (m, 4H, $H_{14}$), 2.210 (t, 2H, $J$ = 7.0 Hz, $H_{28}$), 1.981 (s, 12H, $H_{11}$ and $H_{12}$), 1.900 (m, 4H, $H_{25}$ and $H_4$), 1.739 (t, 2H, $J$ = 7.5 Hz, $H_{27}$), 1.536 (m, 2H, $H_{26}$).

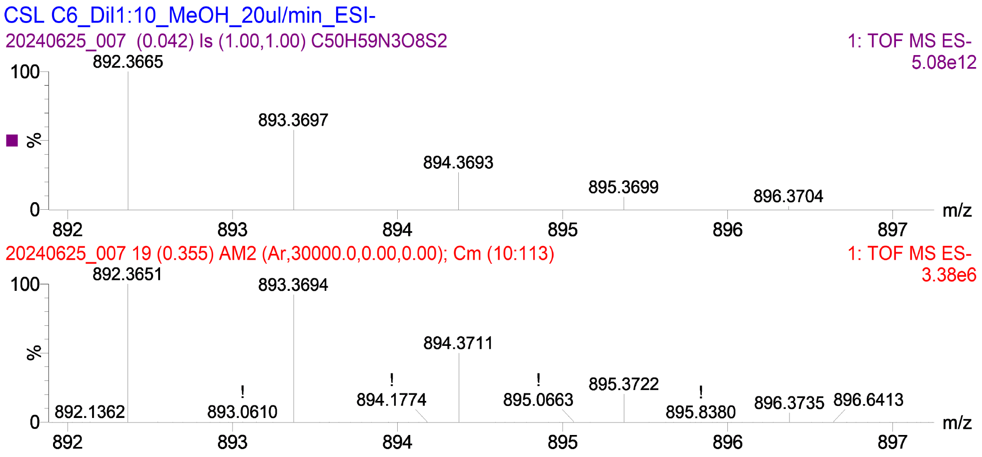


Figure S4: Representative high-resolution ESI mass spectrum of CyC6. Exact-mass values for CyC4, CyC6, and all intermediates are listed with the procedures above.

**HRMS ($\mathbf{ESI^-}$)** calculated for $C_{50}H_{59}N_3O_8S_2$ $[M-H]^-$: $m/z$ 892.3665; found: $m/z$ 892.3651 (Figure S4, Table S2).

Table S2: High-resolution ESI mass-spectrometry data for CyC6.

| $m/z$ experimental | Formula of M | $m/z$ calculated | Proposed ionic species | Error (mDa) | Error (ppm) |
|---|---|---|---|---|---|
| 892.3651 | $C_{50}H_{59}N_3O_8S_2$ | 892.3665 | $[M-H]^-$ | 1.4 | 2 |
| 445.6790 | | 445.6794 | $[M-2H]^{2-}$ | 0.4 | 1 |

LC–MS-grade reagents were used to prepare the sample solution. The sample was analyzed by direct infusion using a high-resolution mass spectrometer equipped with an electrospray ionization (ESI) source and a quadrupole time-of-flight (Q-TOF) analyzer (Xevo G2S Q-TOF, Waters Corp.). Ion-source parameters were optimized to maximize the signal-to-noise ratio in both $ESI^-$ and $ESI^+$ modes.

### Ground-state and steady-state photophysics

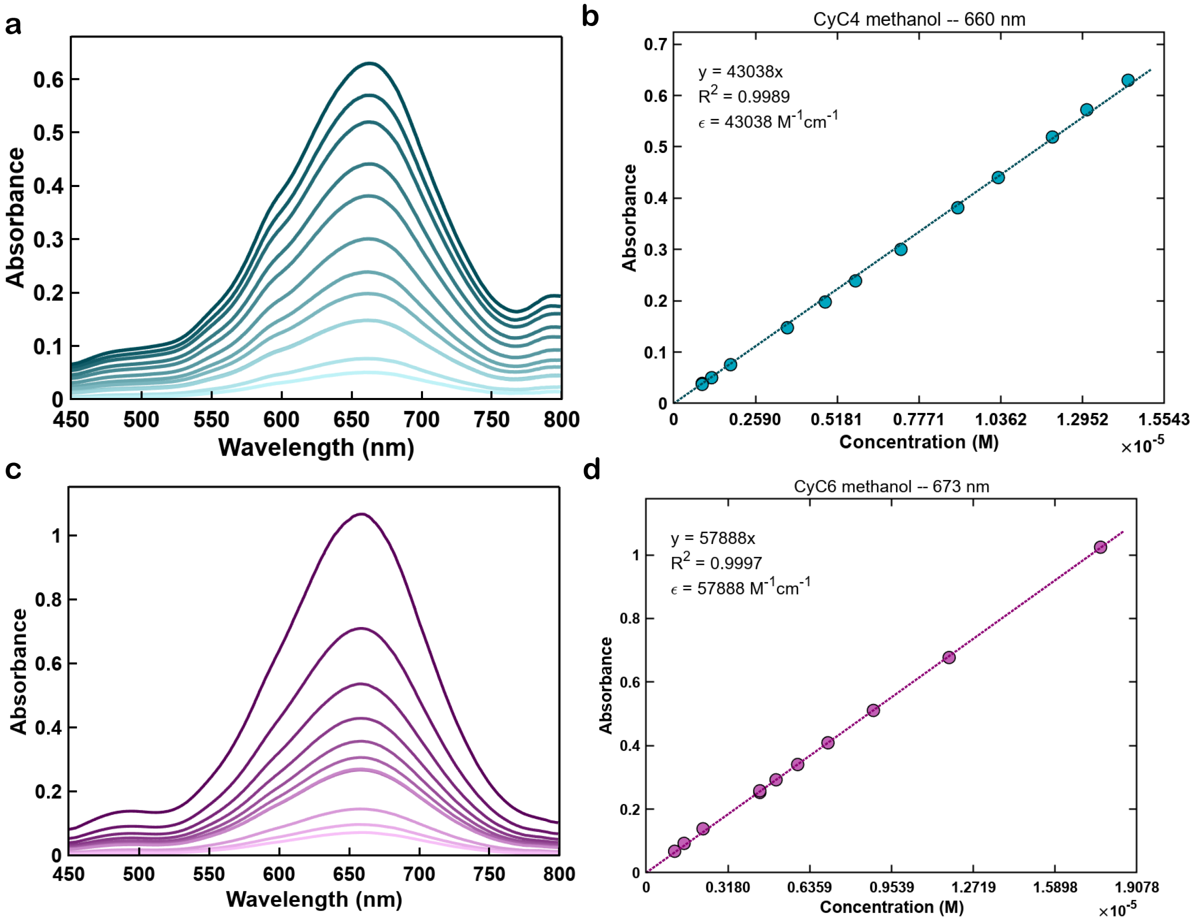


Figure S5: **Molar extinction coefficients of CyC4 and CyC6.** (a) Concentration-dependent absorbance of CyC4 (0.899 –14.4 $\mu$M) in methanol and (b) the corresponding Beer–Lambert fit at 660 nm, $\varepsilon = (4.30 \pm 0.02) \times 10^4$ $M^{-1}$ $cm^{-1}$ (SE, $R^2 = 0.9989$, $n = 14$; two concentrations were measured in technical duplicate). (c,d) The same for CyC6 (1.10–17.7 $\mu$M) in methanol, $\varepsilon = (5.79 \pm 0.02) \times 10^4$ $M^{-1}$ $cm^{-1}$ at 673 nm (SE,$R^2 = 0.9997$, $n = 11$).

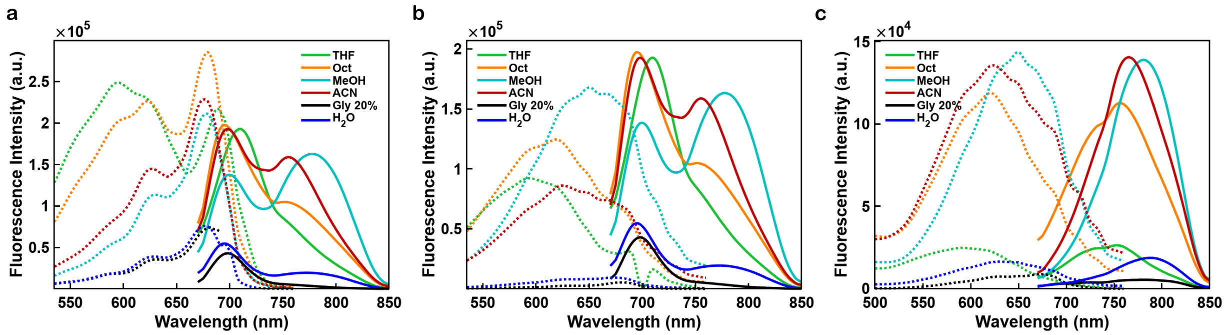


Figure S6: **Overlaid excitation and emission of CyC4 and CyC6 across solvents.** Excitation (dotted) and emission (solid, 660 nm excitation) spectra of CyC4 (0.569 $\mu$M) and CyC6 (0.246 $\mu$M) in tetrahydrofuran (THF), 1-octanol, methanol, acetonitrile, 20% glycerol, and water. (a) CyC4, excitation spectra monitored at 700 nm emission. (b) CyC4, excitation spectra monitored at 780 nm emission. (c) CyC6, excitation spectra monitored at 780 nm emission.

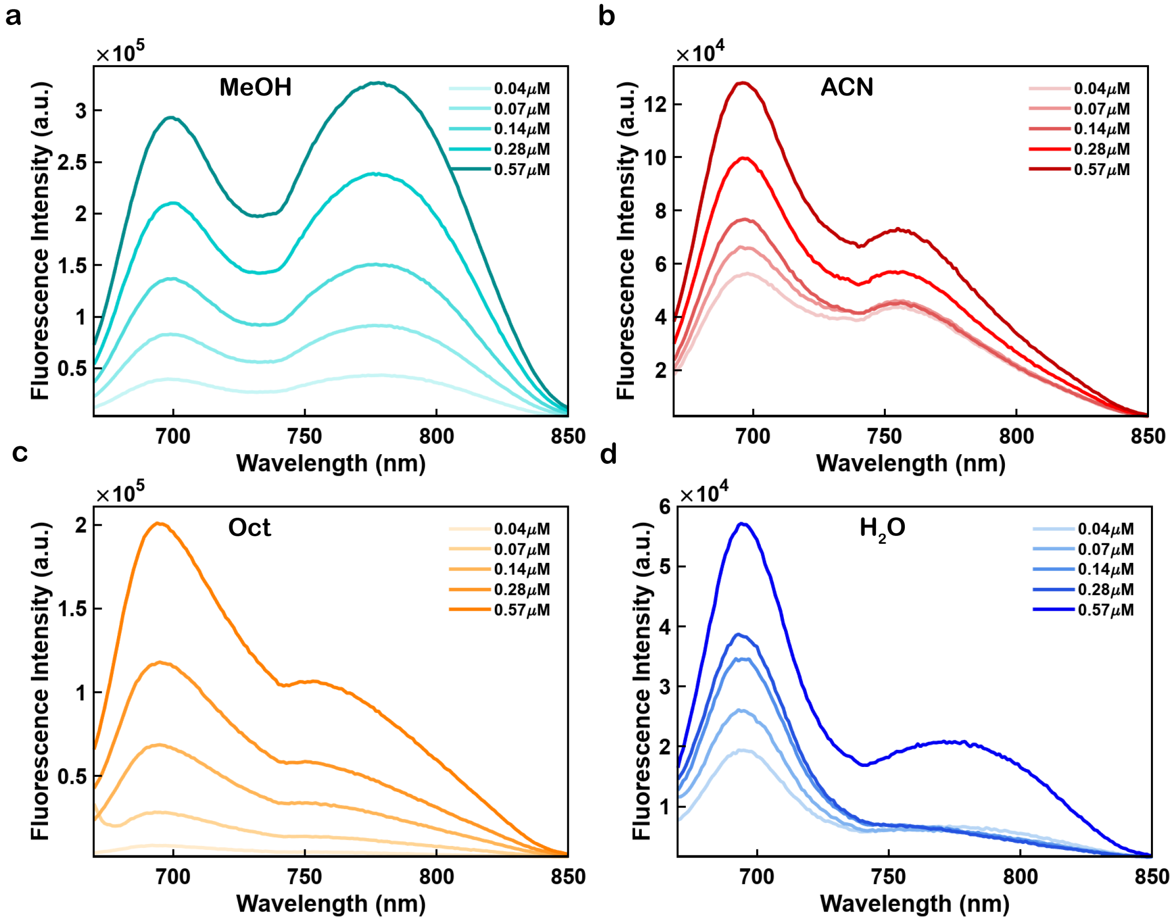


Figure S7: **Concentration-dependent emission of CyC4.** Emission spectra (660 nm excitation) of CyC4 at 0.0357, 0.0711, 0.142, 0.284, and 0.569 µM (rounded to two decimal places in the figure legend) in (a) methanol, (b) acetonitrile, (c) 1-octanol, and (d) water, showing the concentration-dependent growth of the 700 and 780 nm monomer bands with the 700/780 band ratio approximately preserved.

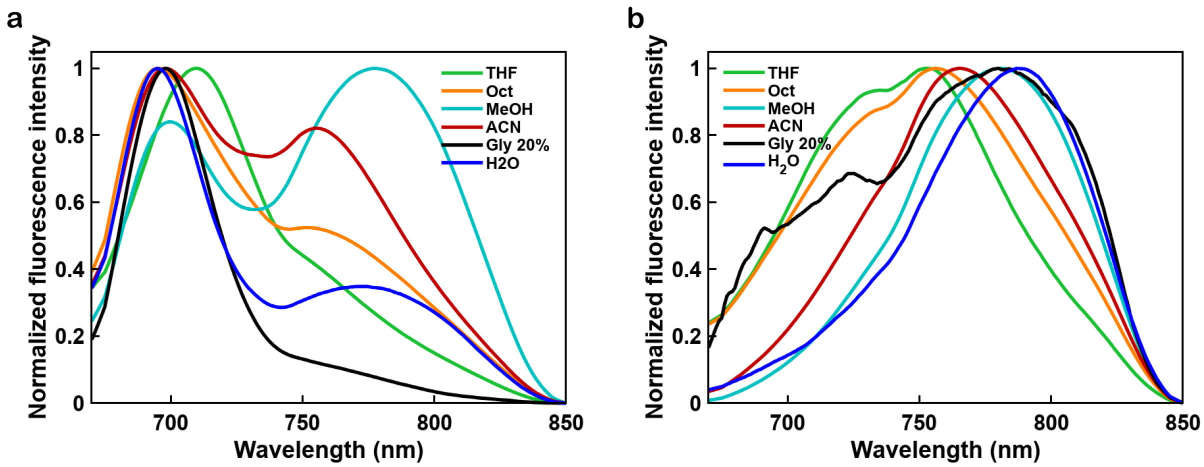


Figure S8: **Peak-normalized emission of CyC4 and CyC6 across solvents.** Emission spectra (660 nm excitation) in tetrahydrofuran (THF), 1-octanol, methanol, acetonitrile, 20% glycerol, and water, normalized to the band maximum. (a) CyC4 (0.569 $\mu$M), normalized to the ~700 nm closed-state (C) band. (b) CyC6 (0.246 $\mu$M), normalized to the ~780 nm open-state (O) band, highlighting the solvent-dependent redistribution between the C-state (700 nm) and O-state (780 nm) bands.

91 # Computational details

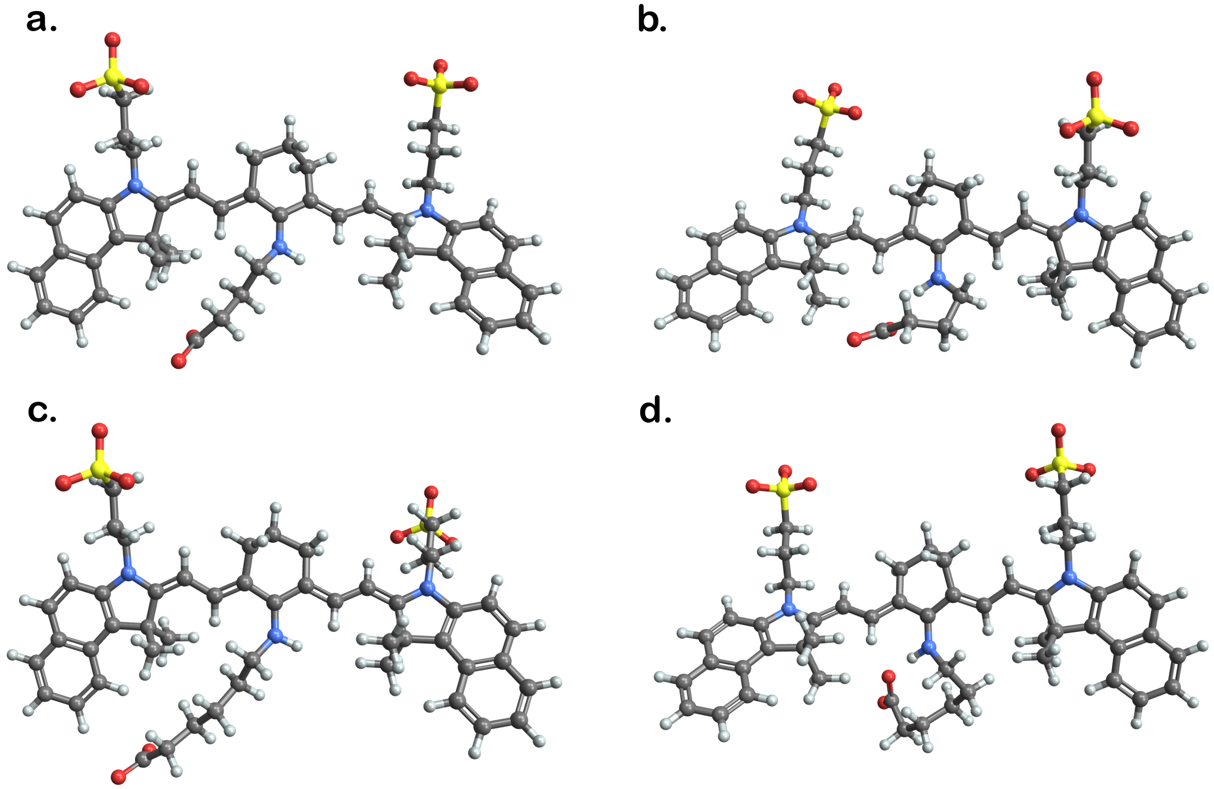


Figure S9: DFT-optimized geometries in methanol ($\omega$B97X-D4/def2-TZVP, CPCM). (a,b) CyC4 conformers; (c,d) CyC6 conformers.

Table S3: Calculated ground- and estimated excited-state intramolecular hydrogen-bond thermochemistry ($-\Delta H_{\mathrm{H-bond}}$, kcal mol$^{-1}$; $\omega$B97X-D4/def2-TZVP, CPCM). Excited-state enthalpies are estimated as the sum of the ground-state enthalpy and the vertical excitation energy calculated by TD-DFT.

| | Water | Methanol | Acetonitrile |
|---|---|---|---|
| CyC4, $S_0$ | -0.15 | 1.95 | 4.04 |
| CyC6, $S_0$ | 1.82 | -1.65 | 4.40 |
| CyC4, $S_1$ | -3.06 | 0.05 | -0.60 |
| CyC6, $S_1$ | 0.01 | -3.19 | 0.48 |

Table S4: TD-DFT vertical excitations and frontier-orbital energies for CyC4 and CyC6 in the non-hydrogen-bonded (O-state) and hydrogen-bonded (C-state) conformers ($\omega$B97X-D4/def2-TZVP, CPCM).

| Dye | Solvent | Conformer | $\lambda_{\max}$ (nm) | $\Delta E$ (eV) | $E_{\mathrm{HOMO}}$ / $E_{\mathrm{LUMO}}$ (eV) |
|---|---|---|---|---|---|
| CyC4 | THF | O | 474.2 | 2.615 | −6.636 / −0.481 |
| | | C | 441.3 | 2.810 | −6.540 / −0.147 |
| | 1-octanol | O | 473.8 | 2.617 | −6.726 / −0.561 |
| | | C | 430.4 | 2.881 | −6.641 / −0.166 |
| | methanol | O | 467.4 | 2.653 | −6.759 / −0.565 |
| | | C | 448.6 | 2.764 | −6.749 / −0.429 |
| | acetonitrile | O | 467.7 | 2.651 | −6.750 / −0.552 |
| | | C | 439.1 | 2.824 | −6.729 / −0.326 |
| | water | O | 472.4 | 2.625 | −6.939 / −0.781 |
| | | C | 450.7 | 2.751 | −6.938 / −0.635 |
| CyC6 | THF | O | 471.3 | 2.631 | −6.665 / −0.491 |
| | | C | 450.9 | 2.750 | −6.570 / −0.235 |
| | 1-octanol | O | 471.3 | 2.631 | −6.755 / −0.573 |
| | | C | 452.1 | 2.743 | −6.670 / −0.338 |
| | methanol | O | 462.2 | 2.683 | −6.779 / −0.539 |
| | | C | 450.9 | 2.750 | −6.754 / −0.427 |
| | acetonitrile | O | 464.4 | 2.670 | −6.763 / −0.537 |
| | | C | 436.8 | 2.839 | −6.744 / −0.302 |
| | water | O | 467.2 | 2.654 | −6.964 / −0.761 |
| | | C | 453.7 | 2.733 | −6.947 / −0.643 |

Optimized Cartesian coordinates and total energies (ORCA, $\omega$B97X-D4/def2-TZVP, CPCM) for the closed (C-state, intramolecularly hydrogen-bonded) and open (O-state) conformers of CyC4 and CyC6 in each solvent are provided in the separate archive `CyC4_CyC6_DFT_optimized_geometries.zip`; each file's comment line reports the total electronic energy in Hartree.

### Aggregation and the crowding channel

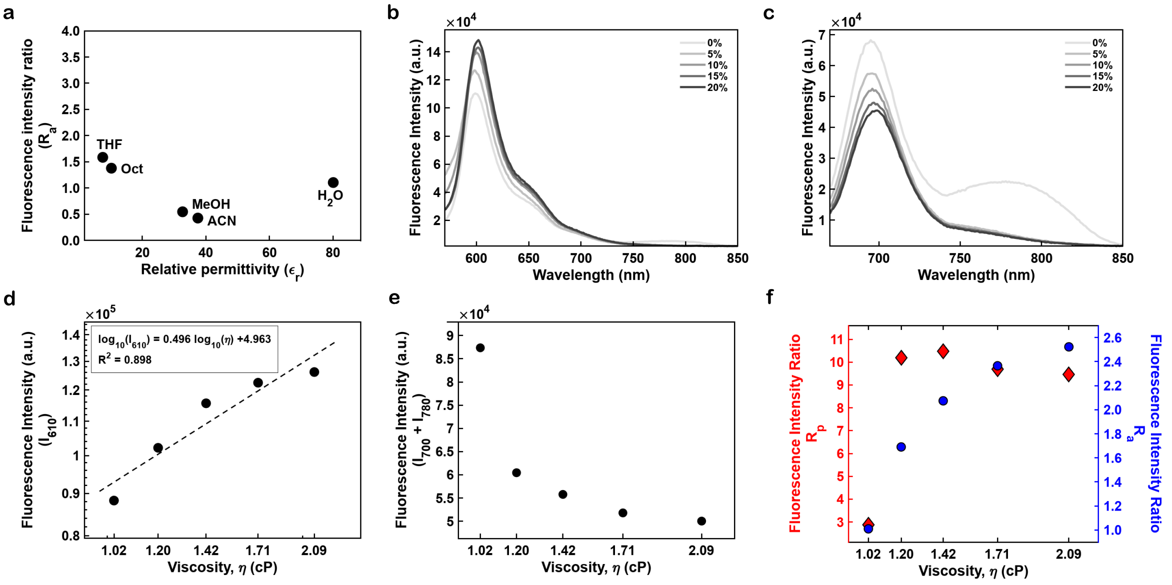


Figure S10: **Solvent- and viscosity-dependence of the aggregation channel.** (a) Aggregation ratio $R_\mathrm{a} = I_{610}/(I_{700} + I_{780})$ versus solvent relative permittivity $\varepsilon_\mathrm{r}$. (b,c) Emission spectra of CyC4 (0.569 $\mu$M) across a glycerol titration (0–20 v/v%) in the ~610 nm and ~700 nm regions. (d) Aggregate-band intensity $I_{610}$ versus viscosity $\eta$ (log–log; $\log_{10} I_{610} = 0.496 \log_{10} \eta + 4.963$, $R^2 = 0.898$). (e) Summed monomer intensity $(I_{700} + I_{780})$ versus viscosity. (f) $R_\mathrm{a}$ (right axis) and the polarity ratio $R_\mathrm{p} = I_{700}/I_{780}$ (left axis) versus viscosity $\eta$: $R_\mathrm{a}$ rises from ~1.0 to ~2.5 while $R_\mathrm{p}$ is essentially constant above $\eta \approx 1.2\,\mathrm{cP}$, establishing $R_\mathrm{a}$ as a reporter of viscosity/self-association and confirming that $R_\mathrm{p}$ is viscosity-independent and largely decoupled from $R_\mathrm{a}$.

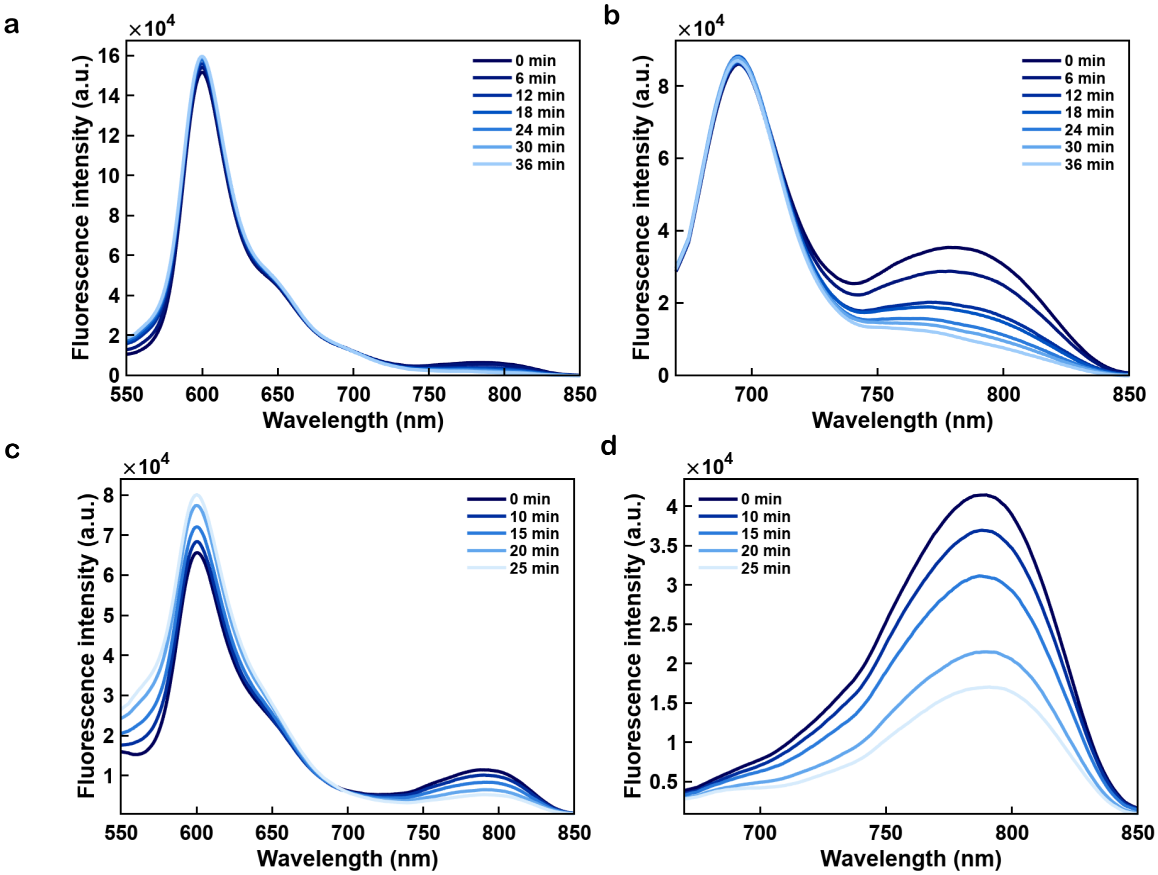


Figure S11: **Time-dependent aggregation of CyC4 and CyC6 in water.** Emission spectra of CyC4 (1.50 $\mu$M, 0–36 min) and CyC6 (0.41 $\mu$M, 0–25 min) in water at room temperature, recorded at fixed time intervals after sample preparation with no additional trigger, using a 0.3 s integration time. (a,c) 540 nm excitation, showing the growth of the ∼610 nm aggregate band over time for CyC4 and CyC6, respectively, consistent with spontaneous self-association. (b,d) 660 nm excitation, showing the corresponding changes in the 700/780 nm bands for CyC4 and CyC6, respectively.
